# Deletion of a conserved genomic region associated with adolescent idiopathic scoliosis leads to vertebral rotation in mice

**DOI:** 10.1101/2023.06.22.546196

**Authors:** Jeremy McCallum-Loudeac, Edward Moody, Georgia Johnstone, Kathy Sircombe, Andrew N. Clarkson, Megan J. Wilson

**Author notes:** corresponding author. ORCID: 0000-0003-3425-5071.

## Abstract

Adolescent idiopathic scoliosis (AIS) is the most common form of scoliosis, in which spinal curvature develops in adolescence, and 90% of patients are female. Scoliosis is a debilitating disease that often requires bracing or surgery in severe cases. AIS affects 2-5.2 percent of the population; however, the biological origin of the disease remains poorly understood. In this study, we aimed to determine the function of a highly conserved genomic region previously linked to AIS using a mouse model generated by CRISPR-CAS9 gene editing to knockout this area of the genome to better understand the biological cause of AIS, which we named AIS_CRM**Λ.** We also investigated the upstream factors that regulate the activity of this enhancer *in vivo*, whether the spatial expression of the LBX1 protein would change with the loss of AIS-CRM function, and whether any phenotype would arise after deletion of this region. We found a significant increase in mRNA expression in the developing neural tube at E10.5, and E12.5, for not only *Lbx1* but also other neighboring genes. Adult knockout mice showed vertebral rotation and proprioceptive deficits, also observed in human AIS patients. In conclusion, our study sheds light on the elusive biological origins of AIS, by targeting and investigating a highly conserved genomic region linked to AIS in humans. These findings provide valuable insights into the function of the investigated region and contribute to our understanding of the underlying causes of this debilitating disease.

## Background

The most common form of idiopathic scoliosis (IS), which spontaneously arises in individuals without pre-existing comorbidities and primarily affects adolescents, is Adolescent Idiopathic Scoliosis (AIS) (Weinstein et al., 2008). Adolescent-onset scoliosis initially develops between 10 −18 years of age and accounts for 85–90% of IS patients, affecting 2-3% of adolescents and 2-5.2% of the total population (de Souza et al., 2013). In addition, scoliosis is the most common developmental disorder affecting the vertebral column (Wise et al., 2020).

While AIS is equally common in both sexes early in the disease course, extreme sex bias develops as curve progression increases (Grauers et al., 2016; Konieczny et al., 2013). As curves progress beyond 40°, females are affected at 10:1 relative to males, with 90% of severely progressive curves requiring surgical intervention. Thus, AIS appears to be driven by genetic susceptibility to unknown internal and/or extrinsic factors. Intrinsic factors are hypothesised to be hormonal fluctuations during puberty, physical disturbances to stable vertical growth in abnormal muscular and ligament tension, changes in pelvic obliquity and size, particularly in women, and altered dorsal horn/interneuron transmission. Extrinsic factors include environmental factors such as exposure to sunlight, altered vitamin D levels, and nutritional status, which can alter pubertal onset and progression, to name a few (Assaiante et al., 2012; Lenssinck et al., 2005; Wong, 2015).

Research on the inheritance patterns of AIS, particularly in twin studies, led to the conducting of GWAS studies (Fadzan and Bettany-Saltikov, 2017; Grauers et al., 2012; Simony et al., 2016). Various genes, including LBX1, have been linked to AIS. LBX1 is a transcription factor that is highly conserved and plays a key role in the development of spinal cord, heart, and limb muscles (Fadzan and Bettany-Saltikov, 2017; Guo et al., 2016; Liu et al., 2017; Takahashi et al., 2011). The SNP rs11190870 is the most commonly associated variant with an increased risk of AIS. It is situated in a highly conserved enhancer element downstream of *Lbx1* (Chen et al., 2014; Guo et al., 2016). A large percentage of disease-associated SNPs, which may be linked to DNA regulation, are found in the non-coding regions of the genome. The onset of diseases in adults is contributed to by tissue-or development-specific roles (Maurano et al., 2012). Guo et al. (2016) found that the DNA fragment had higher activity when the risk SNP (T) was present. Their findings suggest that the SNP rs11190870 disrupts the normal function of the enhancer element, leading to an upregulation of *LBX1* transcription and, therefore, an increased susceptibility to AIS (Guo et al., 2016).

*Lbx1* was first identified in *Drosophila* and has since been identified in metazoan genomes (Jagla et al., 1995). In mice, *Lbx1* mRNA expression is seen as early as Embryonic day 9.5 (E9.5) in the developing neural tube, with levels peaking between embryonic days 11.5-12.5 and decreasing steadily by day E16.5 (Jagla et al., 1995). *Lbx1* is crucial in cell fate, migration, and post-mitotic determination during neural tube development. Gross et al. (2002) observed that many early-born ventrally migrating cells are Lbx1-derived neurons (Gross et al., 2002). These *Lbx1*+ cells, derived from dI4-6 (Class B) dorsal interneuron progenitors, are born between E10 and 12.5 (Gross et al., 2002; Kruger et al., 2002; Muller et al., 2002), post-mitotically expressing *Lbx1* and migrating to populate various regions of the future spinal cord (Muller et al., 2002). In mice lacking *Lbx1*, the substantia gelatinosa within the dorsal horn fails to form, leading to changes in the shape of the grey matter (Gross et al., 2002; Muller et al., 2002). A second Lbx1-positive lineage, born between E11 and E13, migrates dorsally to the substantia gelatinosa of the dorsal horn and other superficial aspects of the dorsal horn (Gross et al., 2002; Muller et al., 2002). These late-born cells are called dorsal interneurons late (dIL) A and B (dIL^A^ and dIL^B^). Altogether, LBX1 plays a crucial role in developing interneuron populations for the proprioceptive network.

The proprioceptive system is a subconscious sensory system that allows individuals to perceive and be aware of their body position in space. This system gathers information on fixed body positions through specialized receptors in the muscles, tendons, joint capsules, and skin (Proske and Gandevia, 2012). In contrast, the kinesthetic system provides information on joint and limb movement. The proprioceptive system functions through a series of reflex arcs found in the spinal cord, which integrate sensory information and help produce smooth and coordinated movement (Dietz, 2002). While the role of the proprioceptive system in controlling posture is well-understood, recent research has directly implicated proprioceptor dysfunction in maintaining spinal alignment (Allum et al., 1998; Blecher et al., 2017). In peripubertal mice lacking TrkC neurons connecting proprioceptive mechanoreceptors and the spinal cord, spinal curvature development was observed without skeletal dysplasia or muscular asymmetry (Blecher et al., 2017).

Studies have shown that patients with adolescent idiopathic scoliosis (AIS) have a deficit in their proprioceptive system, as observed through functional testing of their peripheral joints, such as the knee and elbow (Kouwenhoven and Castelein, 2008; Pialasse et al., 2015). However, this dysfunction’s exact location and timing within the neuronal circuitry remains unknown. Maintaining an upright position requires the integration of the somatosensory, visual, and vestibular systems, with most weighting (∼70%) on the somatosensory system and the remaining on the vestibular system and some on visual input; however, weighting may change as environmental situations change (Peterka, 2002). The reliance on somatosensory input indicates a vital role in proprioception for maintaining spinal alignment and balance, and defects in this system have been linked to AIS (Assaiante et al., 2012; Blecher et al., 2018; Lau et al., 2022; Le Berre et al., 2017). Furthermore, unlike vestibular deficits associated with AIS, proprioceptive deficiencies have been shown to precede vertebral curvature and rotation in animal models (Blecher et al., 2017).

The SNP rs11190870 is located in a putative regulatory region, the adolescent idiopathic scoliosis-associated cis-regulatory module (AIS-CRM), a region highly conserved between mice and humans. To better understand the biological cause of AIS, we generated a mouse model using CRISPR-CAS9 gene editing to delete this region of the genome. Our goal was to determine if the deletion of AIS-CRM affects the expression of nearby genes, especially *Lbx1*. We also investigated the factors that regulate chromatin structure in vivo and whether there are any changes in the spatial expression of LBX1 protein or any phenotypic changes after the deletion of AIS-CRM.

## Results

### Epigenetic analysis of the AIS_CRM genome region

The SNP rs1190870 is located within a highly conserved region near the candidate CRM in human data (Gao et al., 2013; Jiang et al., 2019; Londono et al., 2014; Takahashi et al., 2011). According to variant analyses in human populations, the T allele (considered high risk) is more prevalent than the C allele, with a ratio of approximately 0.59 (T) to 0.41 (C) in most populations (Table 1). This region is located about 10 kb away from the transcriptional start site (TSS) for *LBX1*, situated at the edge of a highly conserved genome region (Fig. 1A).

**Figure 1.**
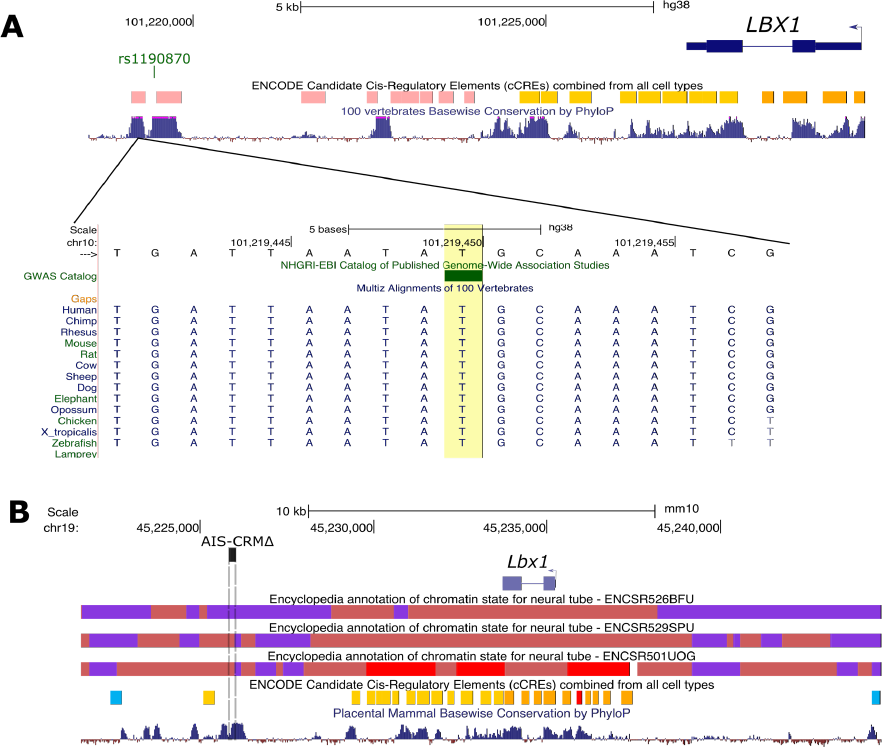
ENCODE regulation data showing candidate CREs and chromatin states for mouse and human *LBX1* loci. **A**. Human *LBX1* gene locus (hg38). The AIS-risk SNP of interest is indicated by green (rs11190870). The ENCODE registry of candidate cis-regulatory elements (cCREs), based on histone modification, DNase, and CTCF binding data, is shown with the corresponding colour key. PhyloP score track based on the alignment of 100 vertebrate species to indicate the level of conservation between vertebrates. **B**. Mouse *Lbx1* locus (mm10 assembly). Chromatin state data (chromHMM 18 state model) for the neural tube (E11.5, E12.5, and E15.5) are displayed along with the colour key. The deleted region in this study is shown (AIS-CRMΔ by the dashed box). PhyloP score track is based on the alignment of placental mammals.

**Table 1.**
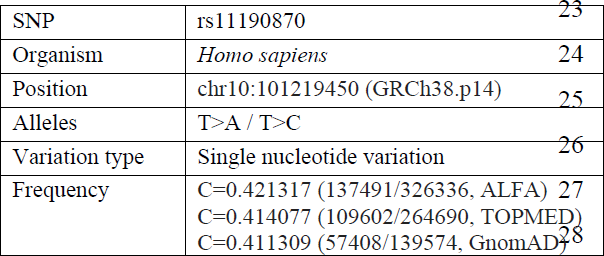
Reference SNP report for rs11190870. Abbreviations: SNP = Single nucleotide polymorph. Data were sourced from (NCBI, 2022)

We analyzed the ChromHMM data for the mouse genome (ENCODE phase 3 data) in the developing neural tube, focusing on the *Lbx1* locus. We found that this region of the genome, which is rich in histone marks, exhibits a bivalent transcriptional start site (TSSBiv) state interspersed between regions repressed by polycomb complex chromatin (PRC2) (Fig. 1B; Indian red). TSSBiv is frequently associated with tissue-specific transcription factor genes and is often assigned to areas with PRC2-bound silencers (van der Velde et al., 2021). We observed similar chromatin landscapes at E11.5 and E12.5. Still, by E15.5, this region shifted to a TSS state (indicating the removal of the repressive mark H3K27me3), suggesting a loss of repressive marks over time (Fig. 1B). This change could potentially affect the expression of nearby genes, such as *Lbx1*. The region has multiple candidate proximal and distal cCREs, most located near the *Lbx1* gene. CTCF-only cCREs are located at each end, with high CTCF signals and low H3K4me3 and H3K27ac candidate sites for chromatin looping or insulators. Although the mouse genomic region corresponding to the AIS-linked SNP in humans is not predicted to lie in cCREs (based on currently available datasets), it lies in a region that shifts from a repressed to a bivalent state, indicating the loss of repressive histone marks (Fig. 1B).

We used RegulomeDB (Boyle et al., 2012) to annotate the SNP-associated region. This database employs various public databases and literature sources to annotate SNP regions. In 123 different tissues, the region that contains rs1190870 was marked as a repressed polycomb. The tissues include "spinal cord", "embryo", "brain", and "gonad" (for the complete list, see File S1). RegulomeDB’s chromatin immunoprecipitation data revealed that enhancer of zeste homologue 2 (EZH2), SUZ12, and RNF2 significantly bind to 10 cell lines derived from a variety of tissues, including embryos (H1 cell line), the spinal cord (neuronal cells), and blood (B cell) (File S1). These proteins are part of the PRC2.

EZH2 is responsible for H3K27 trimethylation and works with multiple TFs to repress gene expression (Gan et al., 2018). To determine whether EZH2 binds to E12.5 brain and neural tube tissues in vivo, ChIP-qPCR was utilized since the RegulomeDB data above was generated with human tissues (Fig. 2B). These tissues were selected because *Lbx1* mRNA is expressed in the neural tube but is barely detectable in the developing brain (Fig. 2C, p = 0.005), which suggests its repression in the CNS of the brain. The AIS-CRM and *Lbx1* TSS areas were targeted. In the brain, EZH2 was significantly enriched in both the AIS-CRM region (Fig. 2B, p = 0.0011) and TSS (Fig. 2B, p = 0.008). However, no significant enrichment was observed in the neural tube compared to the mock control (Fig. 2B, p = 0.096; p = 0.081). In addition, there was significantly more pulldown at these sites within the E12.5 brain than in the neural tube (Fig. 2B, p = 0.0042 and p = 0.0045, respectively).

**Figure 2.**
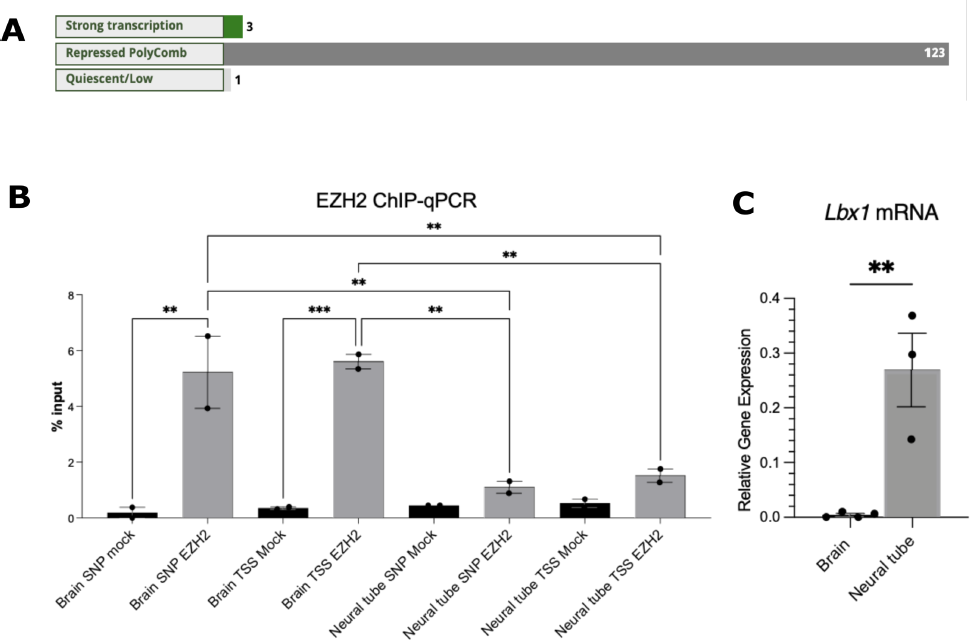
EZH2 targets the *Lbx1* locus for repression. **A**. RegulomeDB (2.0) analysis for chr10:102979206-102979207 found that this region is in a repressed chromatin state for most cell types and tissues. **B**. ChIP-qPCR for EZH2 (n=2). One-way ANOVA with Tukey’s multiple comparison test. ***p<0.001, ** p<0.01. Expression of *Lbx1* mRNA was higher in the neural tube at E12.5, compared to that in the developing brain (n=3-4), two-tailed unpaired t-test.

Together, these epigenetic data suggest that the target region of the mouse genome, which corresponds to the area containing rs1190870 in the human genome, is within the region of the PRC2-bound repressed chromatin.

### Gene expression changes in the AIS-CRMΔ mouse line

AIS-CRMΔ mice were created using CRISPR-Cas9 genome-editing technology, in which RNA-guided nucleases can edit insertions and deletions into specific sites within a genome (Lau and Suh, 2018). The AIS-CRMΔ mouse line removes 189 bp in the mouse genomic region corresponding to the AIS risk SNP rs11190870 (Fig. 1B and 3A). The AIS-CRMΔ adult mice were found to breed normally, suggesting unaffected fertility, and the offspring were viable and displayed typical sex ratios and litter sizes (data not shown).

**Figure 3.**
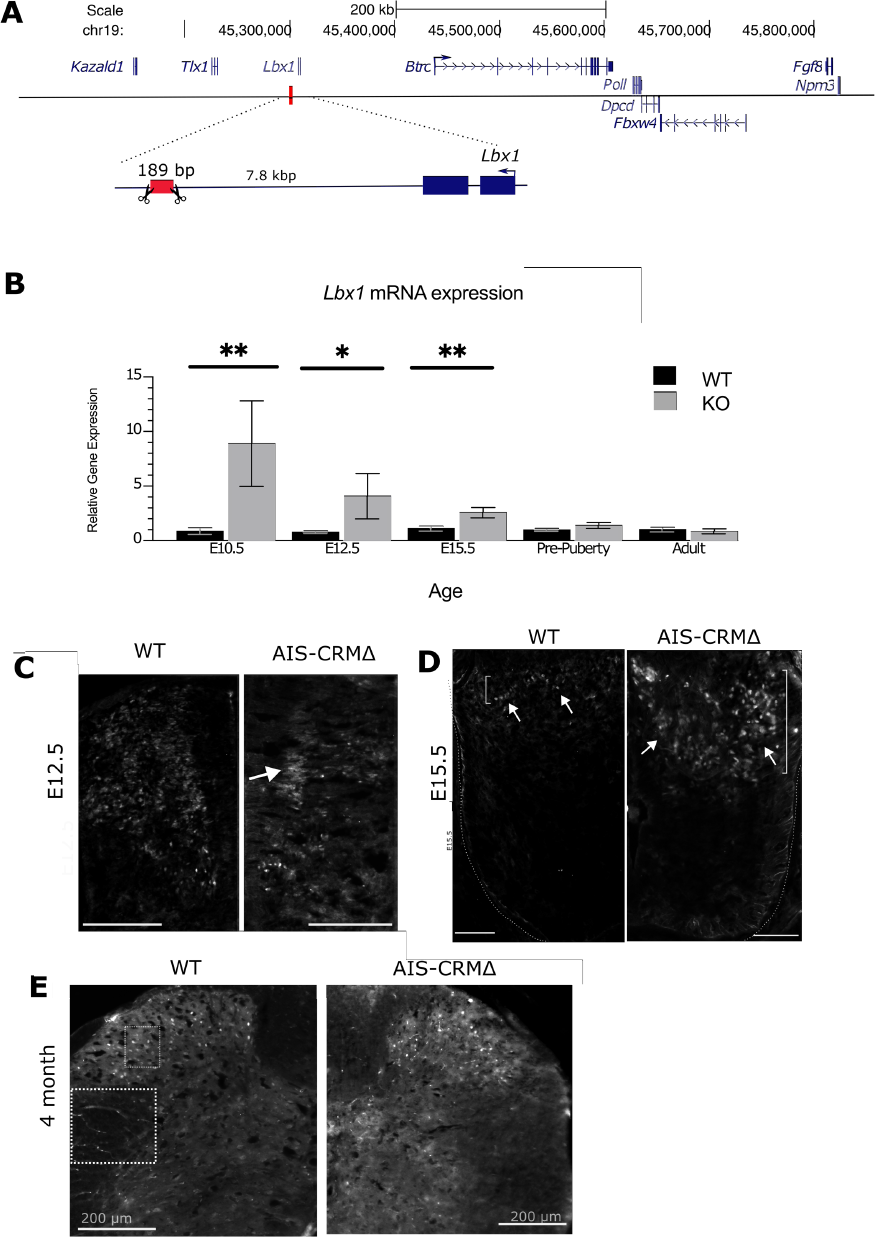
*Lbx1* mRNA and protein expression in WT and AIS-CRMΔ mice. **A.** Schema showing the location of the deleted region in the mouse genome. The red box indicates the deleted region of the mouse model. **B.** RT-qPCR analysis for *Lbx1* transcripts reveals a significant increase in mRNA expression at E10.5, E12.5 and E15.5 in the AIS-CRMΔ embryos relative to WT counterparts. The data is presented as the 2-ΔΔCT relative to WT (mean +/-SEM), and statistical significance was determined using an unpaired t-test (*p<0.05, **p<0.001). The sample sizes for each age group are as follows: E10.5 n=10 (WT), n=3 (AIS-CRMΔ); E12.5 n=11 (WT), n=8 (AIS-CRMΔ); E15.5 n=17 (WT), n=7 (AIS-CRMΔ); Pre-puberty n=9 (WT), n=8 (AIS-CRMΔ); Adult n=6(WT), n=7 (AIS-CRMΔ). (**C-E**) Spatial expression of LBX1 in the neural tube and spinal cord of WT and KO mice. **C.** Representative image of LBX1 protein in the developing neural tube of embryos at E12.5. LBX1 was evident in the marginal (MZ, white arrows) and ventricular zones (VZ, white arrowheads). Expression of LBX1 protein in AIS-CRM embryo neural tube. LBX1 appears less widely spread through the marginal zone (white arrowheads), with increased expression in the ventral region of the neural tube (white arrows) and a few stained cells along the dorsal aspect of the NT. **D**. E15.5 dorsal horn for WT and KO mice **E**. Adult (4 months) spine for WT and KO mice. Scale bar = 100 μm unless otherwise indicated.

Next, we sought to determine how deletion of this putative regulatory region influences gene expression in the neural tube and spinal cord, as this genomic region is predicted to alter the expression of nearby genes. Tissue was collected from three embryonic time points (E10.5, 12.5, and 15.5), representing critical time points during embryonic development, where LBX1 is known to affect neuronal migration and identity (Cheng et al., 2005; Gross et al., 2002; Muller et al., 2002). Additionally, spinal cord mRNA from two postnatal time points (PN28 pre-puberty and PN120 mature) was used to assess changes in the spinal cord before puberty and at maturity.

First, we focused on *Lbx1* mRNA expression, as this gene is predicted to have altered expression *in vivo* because it is located near the target genome region (Fig. 3A). Total RNA was isolated from either pre-(neural tube) or post-natal (spinal cord) time points for WT and AIS-CRMΔ littermates for use in RT-qPCR. We observed a significant increase in *Lbx1* mRNA levels in the AIS-CRMΔ mice at E10.5 (∼9-fold; p = 0.0024) and at E12.5 (5-fold; p = 0.033) and E15.5 (2.3-fold; p = 0.0024) (Fig. 3B. There was no difference in *Lbx1* mRNA expression between WT and AIS-CRMΔ for the pre-puberty and adult spinal cord (Fig. 3B; p = 0.13, p = 0.45).

Following the observation of increased *Lbx1* mRNA expression in the embryonic AIS-CRMΔ neural tube, we assessed the spatial distribution of LBX1 protein. Immunofluorescence experiments were performed on transverse sections of E12.5, E15.5, and adult thoracic samples (Fig. 3C-E). Our results showed that LBX1 protein was present in both the ventricular and marginal zones of the neural tube in E12.5 WT mice (Fig. 3C), consistent with previous studies. In E12.5 AIS-CRM samples, LBX1 protein was also present in both zones (Fig. 3C). In E15.5 mice, LBX1 protein was found in a specific area of the dorsal horn, and in KO mice, the expression was slightly expanded within the substantia gelatinosa of the dorsal horn. Additionally, scattered LBX1-positive cells were found in Lamina II/III and white matter oligodendrocytes in adult mice. Our findings were confirmed by single-cell RNA-seq data (Franzen et al., 2019), which showed LBX1 expression in spinal cord neurons and a small number of oligodendrocytes in adult spinal cords (Fig. S1). These results are consistent with previous studies on LBX1 expression (Gross et al., 2002).

### AIS-CRM is located at the boundary of the TAD

Additionally, we also delved into whether the AIS-CRM knockout affected the expression of nearby genes. The 3D structure of chromatin and distal regulatory elements allow them to interact with genes across long genomic distances. The genome is organized into topologically associated domains (TADs) within the nucleus. These domains are enriched in intradomain chromatin interactions, isolated from surrounding chromatin and bounded by narrow segments known as boundary regions, appearing as triangles (as shown in Fig 4). TADs are highly conserved and play crucial roles in determining cell fate during development (Dixon et al., 2012; Li et al., 2022).

**Figure 4.**
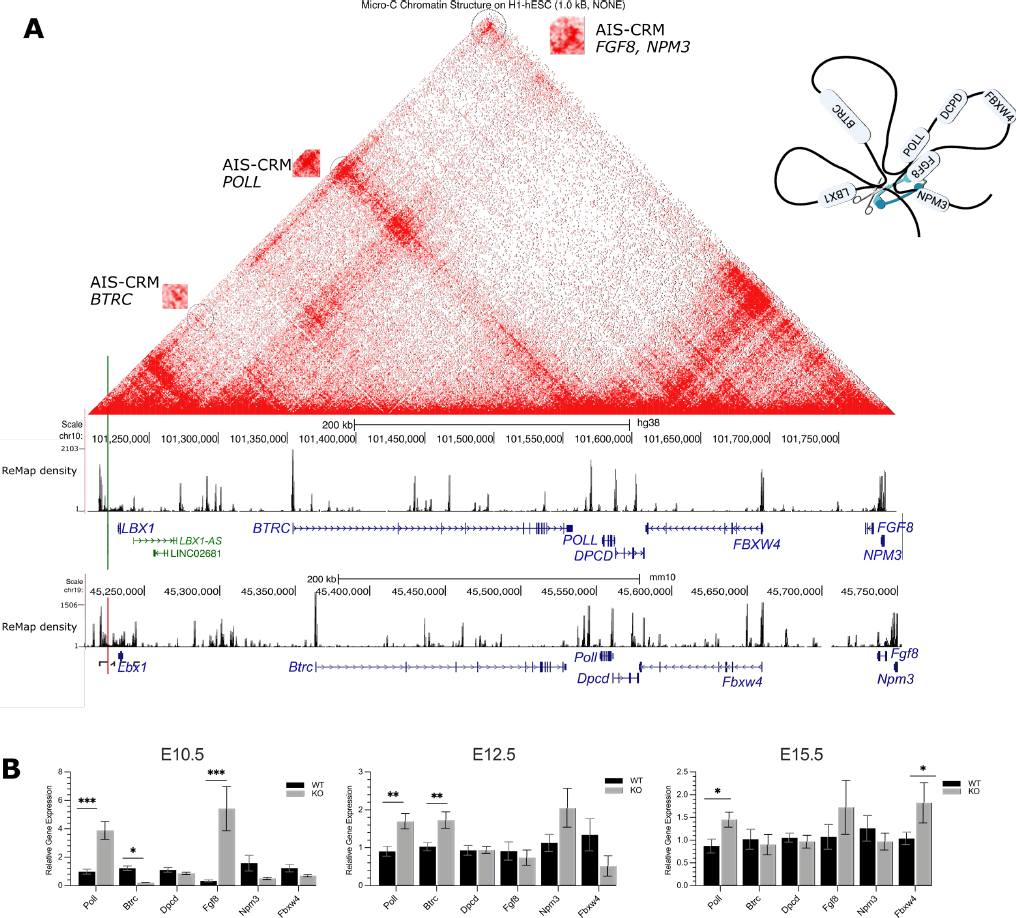
Transcript expression of genes predicted to be in physical contact with the AIS-associated regulatory region *in vivo*. **A.** Human genome (hg38) browser, with micro-C data from human embryonic stem cells (ESC) shown at 1.0 kb resolution and ReMap data. The green vertical line indicates the position of rs1119870 SNP. The red line indicates the deleted region of AIS-CRM. The contact points are circled and enlarged. The density track (ReMap) shows the density of overlapping ChIP-seq peaks. These regions were enriched with CTCF peak regions. In both mice and humans. TAD and internal sub-TAD. The red colour range indicates the interaction frequency (0=none, red=high). **B.** The expression of *Poll* and *Fgf8* mRNA significantly increased in the AIS-CRMΔ group compared to the WT group during the neural tube development at E10.5 (p < 0.001). At E12.5, the AIS-CRMΔ cohort showed approximately 1.8-fold higher levels of *Poll* and *Btrc* transcripts than the WT group (p < 0.05). Similarly, at E15.5, there were significant increases in the expression levels of *Poll* and *Fbx4* mRNA (p < 0.05). These results are presented as mean ±SEM. The statistical significance was determined using two-way ANOVA, and it is presented as *p<0.05, ***p<0.001. The sample sizes for each group were as follows: E10.5 n = 10 (WT), n = 3 (AIS-CRMΔ); E12.5 n = 11 (WT), n = 8 (AIS-CRMΔ); E15.5 n = 17(WT), n = 7 (AIS-CRMΔ); pre-puberty n = 9 (WT), n = 8 AIS-CRMΔ); adult n = 7 (WT), n = 6 (AIS-CRMΔ).

After analyzing available HiC and Micro-C chromatin structure data, we discovered potential contacts and a local TAD structure (Wang et al., 2018) (Fig. 4A and Fig. S2). LBX1 and six other genes were identified within a single TAD, including Fibroblast growth factor 8 (*FGF8*), F-box and WD-repeat domain containing 4 (*FBXW4*), β-transducin repeat containing e3 ubiquitin-protein ligase (*BTRC*), deleted in primary ciliary dyskinesia (*DPCD*), and DNA polymerase lambda (*POLL)*. Additionally, there are sub/internal TAD interactions, such as the contact between AIS-CRM/TAD boundary and POLL and BTRC.

Thus, the rs11190870 variant is located at the TAD boundary, which was also confirmed by ReMap data. This extensive ChIP-seq database can be accessed through the UCSC browser (Hammal et al., 2022). The data shows several dense peak areas, including a similar pattern between mouse and human genomes (Fig. 4), regions near the SNP enriched with CTCF, cohesion proteins (SMC3, RAD21), and Polycomb complex (CBX7, EZH2, JARID2) proteins (zoomed-in view, see Fig. S3). Cohesin subunit proteins and CTCF overlap at boundary regions (Zuin et al., 2014). PRC proteins (EZH2, CBX7, and JARID7) are also associated with genome organization and chromatin looping (Kheradmand Kia et al., 2009). They are enriched here, consistent with the previous results (Fig. S3 and Fig. 2).

In a similar manner to *Lbx1*, the RT-qPCR data showed a significant increase in *Poll* and *Fgf8* mRNA expression at E10.5 in AIS-CRMΔ mice (Fig. 4B). The increase was four-fold for *Poll* and 16-fold for *Fgf8*, respectively (Fig. 4B). *Btrc* mRNA expression decreased by approximately 5.9-fold in AIS-CRMΔ embryos. At E12.5, *Poll* and *Btrc* mRNA levels were approximately 1.8 times higher in the AIS-CRMΔ group compared to the WT group (Fig. 4B). *Poll* mRNA levels remained higher than WT at E15.5, ∼1.7-fold (Fig. 4B). Additionally, the expression of *Fbxw4* mRNA was significantly increased at this stage, ∼1.8-fold (Fig. 4B).

There were no significant differences in the expression of any of the tested genes between WT and KO mice within the pre/post-puberty spinal cords (Fig. S4; p > 0.05). In addition, we also tested a gene just outside the TAD region, Nucleophosmin/nucleoplasmin 3 (*Npm3*); this gene showed no change in mRNA expression (Fig. 4B). Together, these results suggest that the AIS-CRM region of the mouse genome can influence the expression of other genes located within the same TAD during development.

### Phenotypic changes to the vertebral column

We hypothesised that any morphological abnormalities of the spine (vertebral column) would be subtle, partly because human patients with AIS are healthy until adolescence and because of the limited scoliotic curves observed in quadrupeds (Bobyn et al., 2015; Gorman and Breden, 2009). In addition, rodents are known for their inability to reproduce lateral/3-dimensional rotational curvatures unless under severe perturbation, such as the knockout of a critical developmental gene (Adham et al., 2005; Blecher et al., 2017; Sparrow et al., 2013). Therefore, we examined the gross morphology of the vertebrae using micro-CT scanning, followed by analysis of vertebral rotation.

To assess gross spine morphology, the reconstructed microCT scans were examined using the 3D Viewer plugin for FIJI. No apparent differences in the shape or profile of the vertebral bodies, pedicles/lamina, vertebral foramen, or spinous processes were observed (Fig. S5). We could not find obvious evidence of ultrastructural changes to the vertebrae at 34 μm resolution (data not shown). Vertebral rotation is a critical feature of scoliosis, and the spinal curvature indicates vertebral rotation (Janssen et al., 2010; Lam et al., 2008; Lee et al., 2020; Xiong et al., 1993). Therefore, we determined the vertebral rotation in our mouse line. To ensure accurate and comparable results, the 5^th^ Lumbar vertebrae (L5) was selected as the ‘anchor’ with its rotation set as 0°, allowing us to measure vertebral rotation relative to L5 in all samples.

We noted considerable variation in vertebral rotation between the genotypes (Fig. 5). However, the results showed that the variation between the two groups (over the entire vertebral column) was not significantly different and that pre-puberty WT and AIS-CRMΔ mice exhibited similar variation in rotation of the vertebrae (Fig. 5; p = 0.43). When we examined variation at the individual vertebral level, that is, comparing the variation in standard errors of the mean between WT and AIS-CRMΔ mice, we observed a greater propensity for rotation in AIS-CRMΔ mice. Thus, KO individuals had a greater variation than in WT mice.

**Figure 5.**
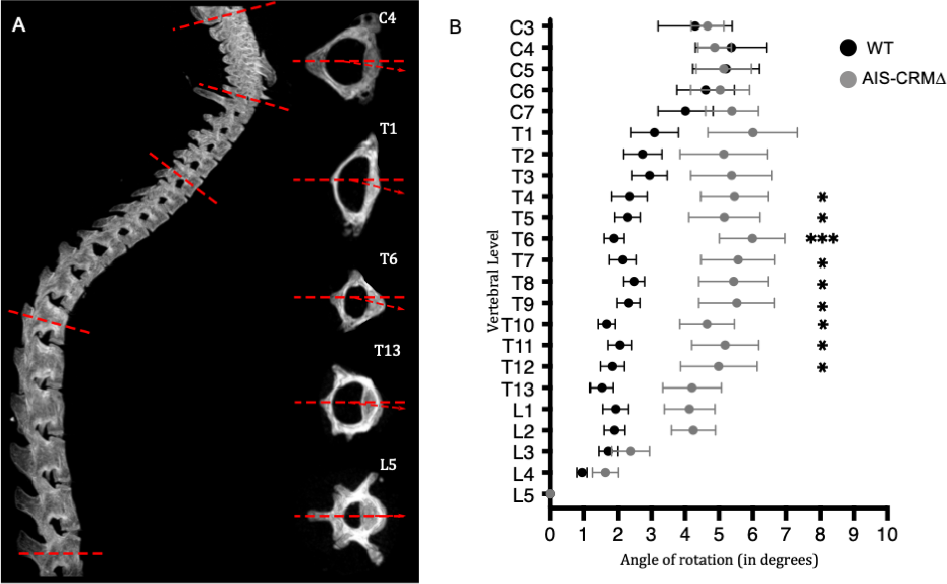
Vertebral rotation in adult WT and AIS-CRMΔ Knockout mice using μCT. **A.** Lateral view of an example spine (left) and the acquisition of rotational measurements using individually reconstructed vertebrae (right). Red dashed line shows baseline, 0° rotation, red arrow shows deviation of that vertebra. B. Degree of rotation measured across C3-L5 in the mouse samples, combining sexes. Wild-type (WT) mouse measurements are represented in black (n = 20) and AIS-CRM**Δ** in gray (n = 10). Measurements are presented as mean rotational deviation; error bars represent ± SEM; statistical significance is represented as p < 0.05 =*, p < 0.01=**, p < 0.001=*** using multiple unpaired t-tests with multiple comparison correction (two-stage step-up (Benjamini, Krieger, and Yekutieli)).

In human AIS, curve apices are found in the central vertebrae of the affected region (i.e., the cervical apex is typically C3-4, thoracic T6-T7, thoracolumbar T12-L1) (Busscher et al., 2010; Lenssinck et al., 2005; Wong, 2015). Thoracic and thoracolumbar curves have been observed in 88% of AIS cases (Greiner, 2002; Schlosser et al., 2014; Weinstein et al., 2008). This led us to focus on the thoracic region in the mouse cohort. We found a significant rotational deviation in the thoracic region, T4-T12, in the AIS-CRM cohort (Fig. 5). Considering rotational variation across all samples, the most considerable degree of rotational deviation is at T6, with a mean of 6° (± 2.94; p < 0.001) (Fig 5). Vertebral rotations for T4–T12 were all significantly more rotated than their WT counterparts (p < 0.05), with a larger variation in AIS-CRM samples than their WT counterparts, as evidenced by the larger SEM (Fig. 5). Subsequently, a two-way ANOVA (pooled) was performed to determine whether the genotype had a similar effect at all vertebral levels, and 4.3% of the variation (P = 0.02) suggested an interaction.

Due to the sex-biased nature of AIS, we sought to determine whether this difference between WT and AIS-CRM was influenced by biological sex (Fig. S6). The largest rotation angle was nearly identical between our male (6.416° ± 3.491; T6) and female cohorts (6.415° ± 2.824; T8) (Fig. S6). We conducted a comprehensive 3-way ANOVA analysis (File S2) to investigate potential interactions among genotype (WT/KO), sex, and vertebrae level. Notably, our findings revealed a significant effect of genotype (p = 0.0001), indicating that the presence or absence of the conserved genomic region associated with AIS in humans plays a pivotal role in vertebral rotation. However, we did not observe any significant interaction between the factors of sex, genotype, and vertebral level. This implies that the impact of the AIS-CRM deletion on vertebral rotation remains consistent regardless of sex.

Vertebral rotation was also analysed in pre-pubertal mice (< PN28). There were no statistically significant differences between the two cohorts (Fig. S7; p = 0.44). These results (Fig. 5 and Fig. S7) suggest that the AIS-CRM mice’s vertebral rotation onset occurred following pubertal onset.

### Simple proprioceptive behaviour was affected in AIS-CRMΔ mice relative to their WT littermates

Having demonstrated rotational instability of the vertebrae in AIS-CRMΔ mice, we sought to determine whether these mice exhibited proprioceptive deficits following the findings of Blecher et al (2017), linking proprioceptive system disruption to scoliotic/vertebral instability (Blecher et al., 2017). This was particularly interesting given the influence of LBX1 on neural tube interneuron development and the potential for the proprioceptive system to be influenced by the altered expression observed above.

Initially, we performed a modified SNAP test (Simple Neuroassessment for asymmetric impairment), as previously described (Shelton et al., 2008), at four time points:4 (pre-puberty) and 16, 20, and 24 weeks of age. We found that across all time points, AIS-CRMΔ mice scored higher than their WT littermates; this difference was statistically significant at 4 (p = 0.0043) and 16 weeks (p = 0.0093) (Fig. 6). The statistically significant increase in the SNAP score from 4 to 16-week (p = 0.012) for the knockout mice coincided with pubertal transition; however, this was not observed in the WT cohort (p = 0.478) (Fig. 6). At 20 weeks of age, WT and AIS-CRM mice performed equally in the SNAP test. This trend continued until the end of testing at 24 weeks of age (P = 0.12). Together, this suggests a possibility of a pre-mature decline in proprioceptive function for AIS-CRMΔ and confirms previously reported changes in age-related proprioceptive functions.

**Figure 6.**
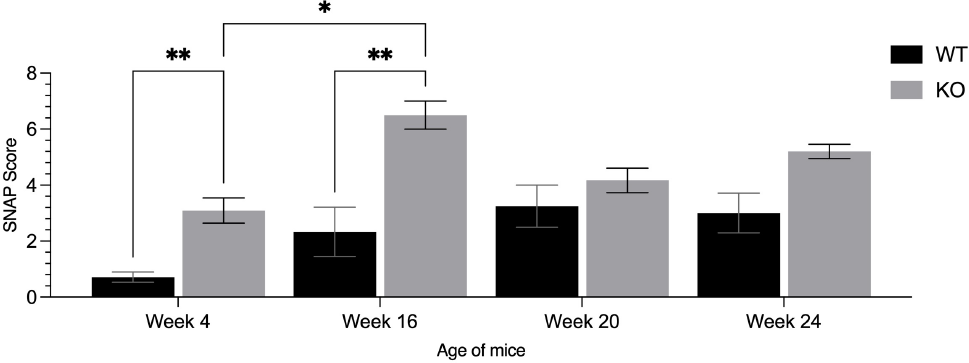
SNAP testing of AIS-CRMΔ revealed premature proprioceptive deficits. The results of the SNAP score suggest that at 4 and 16 weeks of age, AIS-CRMΔ mice performed worse in simple proprioceptive tasks than their WT counterparts (p = 0.0043; p = 0.0124). However, at 20 weeks of age, the mice performed equally in the proprioceptive tests (p > 0.05), which continued until 24 weeks of age. Data are presented as the mean ±SEM. Statistical significance was determined using two-way 2-Way ANOVA where ** p < 0.01. Abbreviations: SNAP, **simple** assessment of **asymmetric** im**P**airment; WT, wild-type; AIS-CRM.

We used a cylinder test to evaluate the CNS circuits in mice that control simple reflexes and motor function (Roome and Vanderluit, 2015). We wanted to know if AIS-CRM mice had problems with basic motor skills, including raising, choosing their paws, controlled paw removal, and other noticeable behaviours such as stumbling or falling (Fig. S8). However, between 8 and 24 weeks, we only noticed a very modest reduction in the time spent rearing in the WT and AIS-CRM groups, which was not statistically significant (Fig. S8; p = 0.28, p = 0.09, respectively).

Although the AIS-CRMΔ mice spent a similar amount of time rearing as mice without the AIS-CRMΔ mutation, they did exhibit some functional impairments in their forelimbs. This was further confirmed through the grid walk and grip strength tests (Figs. 7 and S9). Additionally, when standing and rearing, the AIS-CRM mice exhibited a propensity to fall forwards to the ground because of hindlimb instability (Fig. 7A). Furthermore, their hindlimbs had a wider stance and were more splayed compared to their WT counterparts (Fig. 7B).

**Figure 7.**
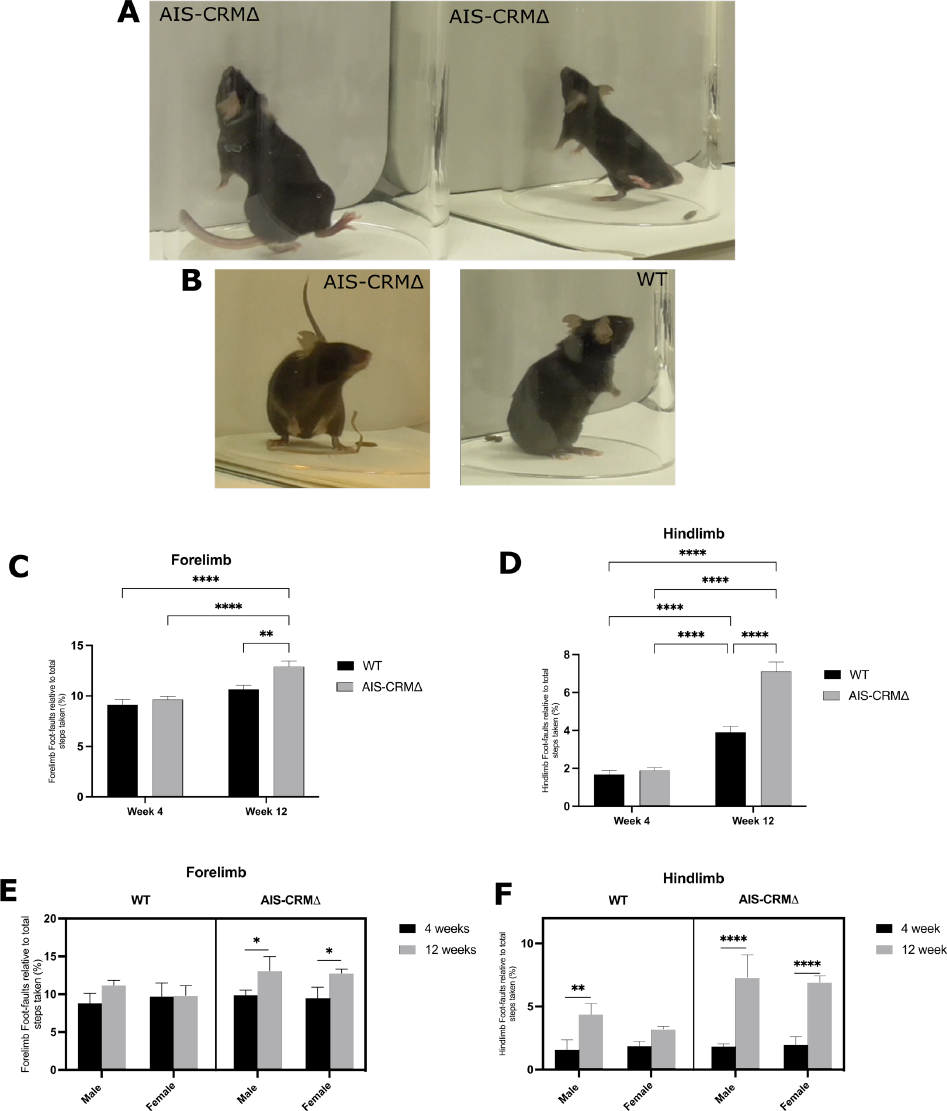
Observational functional deficits were observed in AIS-CRMΔ mice. A) AIS-CRMΔ mice showed a loss of hindlimb balance when reared and falling forward towards the ground after failing to touch the cylinder wall. B) The AIS-CRMΔ mouse (left) was observed with a wider hindlimb stance when reared and on the ground than their WT counterparts (right). Effect of the AIS-CRMΔ deletion on the number of limb foot-faults over 8 weeks. C) Forelimb foot faults relative to the total steps taken for the WT and AIS-CRMΔ. D) Hindlimb foot faults relative to the total steps taken by WT and AIS-CRMΔ. All data are expressed as the mean ± SEM (n = 8 WT and n = 7-13 AIS-CRMΔ), and statistical analysis using two-way ANOVA is represented as *p < 0.05, **p < 0.01, ***p < 0.001, and ****p < 0.0001. E) Forelimb data separated by sex (n = 5 WT and n = 4-5 AIS-CRMΔ). F) Hindlimb foot faults separated by sex (n = 3 WT and n = 3 - 8 AIS-CRMΔ). The data are shown as a three-way plot separated by AIS-CRM genotype, age, and sex. Statistical analysis was performed using three-way ANOVA (mixed effects) and Tukey’s multi-comparison test. Abbreviations: WT, wild-type; AIS-CRM, Δ knockout mice.

The grid walk test evaluates the locomotor function and motor deficits in rodents with CNS disorders (Baskin et al., 2003; Lin et al., 2016; Schaar et al., 2010). It is also commonly used to assess the sensorimotor abilities of the forelimbs and hindlimbs (Russell et al., 2011). Our goal was to determine if the deletion of certain genes affected the rodents’ functional abilities, specifically by measuring the number of foot faults relative to the total number of steps taken per limb (Fig. 7C and D). We found a modest increase in forelimb foot faults in AIS-CRMΔ mice as they aged from 4 to 12 weeks, which was statistically significant. This increase was not observed in their WT litter-mate mice over the same period. Additionally, at 12 weeks, there was a significant difference in the number of forelimb foot faults between the WT and AIS-CRMΔ mice (1.3-fold; p = 0.0087; Fig. 7C). This suggests that the deletion of AIS-CRMΔ results in sensorimotor dysfunction of the forelimb during puberty, which leads to an increase in the number of foot faults.

The WT and AIS-CRMΔ cohorts showed a significant increase in hindlimb foot faults from 4 to 12 weeks of age. The WT cohort had a 2.3-fold increase (p < 0.0001), while the AIS-CRMΔ cohort had a 3.7-fold increase (p < 0.0001). Although both cohorts experienced an increase in hindlimb foot faults, the AIS-CRMΔ mice had a 1.8-fold increase (p < 0.0001, Fig. 7D) in the total number of hindlimb foot faults compared to the WT mice. This indicates that while there is a natural decline in function during this period, the deletion of AIS-CRM leads to a more significant decrease in sensorimotor function.

We conducted a three-way ANOVA analysis using the mixed-effects model REML to examine how sex, AIS-CRM genotype, and age (in weeks) interacted. The results are summarized in File S2. Genotype and age significantly impacted forelimb foot faults (p = 0.048; Fig. 7C), while sex did not affect the outcome (File S2). When we separated the results by sex, male and female AIS-CRMΔ mice showed a significant increase in forelimb foot faults from four to 12 weeks (Table SX). For the hind limb, there was a significant interaction between genotype and age (p = 0.0003), but not with sex. Both sexes observed significant increases in hindlimb foot faults (File S2). Female WT mice showed no significant changes in the number of hindlimb foot faults from week 4 to week 12 (p = 0.57, File S2). However, male WT mice experienced a significant increase in hindlimb foot faults (2.8-fold, P = 0.0047; File S2) over an 8-week period. This suggests that WT male mice may experience a decline in hindlimb locomotor function over time. Male and female AIS-CRMΔ mice also showed a significant increase in hindlimb foot faults over eight weeks (p < 0.0001) (Fig. 7F; File S2). This trend continued at 12 weeks of age, with 1.7- and 2.2-fold increases in male and female AIS-CRMΔ mice, respectively (p<0.0001, Fig. 7F; File S2).

The grip strength test is used to evaluate motor function and deficits in mice with CNS disorders (Maurissen et al., 2003). WT and AIS-CRM mice showed a decline in forelimb grip strength over time, with no significant difference in strength between the two groups (Fig. S9, p = 0.08, p = 0.65, respectively). However, there was a significant decrease in forelimb grip strength in WT mice from 4 to 14 weeks (p = 0.004) and from 4 to 24 weeks (p = 0.002), with a 38% and 47.8% decline in strength, respectively (Fig.S9). These decreases likely reflect the natural decline in motor function with age, with no apparent differences in grip strength between WT and AIS-CRM mice. During grip strength testing, AIS-CRMΔ mice behaved differently from WT mice, struggling to hold onto the bar and twisting and turning. This behavior was consistent with observations during SNAP testing, where AIS-CRMΔ mice repeatedly failed to perform the baton grip task appropriately (Fig. 6).

## Discussion

Here we investigated a genomic region linked to AIS to understand its function. By analyzing data from a mouse model and using bioinformatics and ChIP-qPCR, we discovered that EZH2 binds to this region near the TAD boundary. Deleting this area in mice caused changes in gene expression in the developing neural tube. This affected the nearby Lbx1 gene and other genes within the same TAD. Our analysis of AIS-CRMΔ mice showed that they had significant vertebral rotation and variation compared to WT mice. Before any morphological changes occurred, these mice also displayed proprioceptive deficits and sensorimotor decline. We believe deleting this genomic region disrupts connectivity in the developing spinal cord. This disruption then leads to proprioceptive deficits and spinal misalignment.

### Gene regulation and AIS-CRM

According to previous studies, AIS-CRM regulates *Lbx1* expression during critical stages of embryonic development, contributing to neural tube patterning and migration. An in vitro luciferase reporter assay conducted by Guo et al. (2016) showed that the risk SNP increased luciferase activity and that spinal curvature was caused by overexpression of *lbx1* in zebrafish (Guo et al., 2016). Additionally, we observed an increase in *Lbx1* expression when the region containing the SNP was knocked out. This increase could be attributed to either an increase in the number of cells expressing *Lbx1* and their protein expression or the detection of cells expressing increased levels of *Lbx1*. Gross et al. used both antibody and GFP line, and found more broad expression in the GFP line, which they suggested was more sensitive to pick up *Lbx1* expression (Gross et al., 2002). The observed increase in expression could be due to an increase in the number of cells that normally express lower levels of *Lbx1*, now detectable with the antibody, or an increase in the number of those cells which express *Lbx1* and their relative increase in protein expression. We also found that deletion of this region increased the expression of nearby genes.

We initially hypothesised a regulatory role for this region in the presence of a PRC2 repressor. PRC2-bound silencers are distal regulatory elements that mediate long-range chromatin interactions with target genes, and disruptions such as CRISPR-Cas9 mediated deletion can lead to the activation of the target gene expression (Ngan et al., 2020). In addition, PRC2-bound silencers can transition into enhancers in specific cell lineages (Ngan et al., 2020). Changes in TAD boundaries are associated with changes in gene expression in cell cultures. TAD boundaries restrict the interactions of cis-regulatory elements with genes within the TAD, and the loss of TAD boundaries results in ectopic gene expression in *in vitro* cell cultures (McArthur and Capra, 2021). Furthermore, TAD boundaries often contain housekeeping genes and transcriptional start sites (Dixon et al., 2012).

### Possible consequences of changes to *Lbx1* gene expression

Although we observed significant differences in the expression profiles of WT and AIS-CRMΔ mice during embryonic development, these observations were no longer evident in our adult cohorts. This finding did not suggest any differences in the adult spinal cord. More importantly, as previously indicated, changes in *Lbx1* expression through development likely led to changes in downstream effectors, consequently triggering changes in neuronal wiring, neuron identity, and overall functionality. *Lbx1* has been implicated in the selection of GABAergic cell fate, and loss of *Lbx1* results in glutamatergic neuron fate. Overexpression of *Lbx1* has recently been shown to alter the neuronal population distribution within the neural tube (Decourtye et al., 2022). Neuron numbers may remain similar between AIS-CRMΔ and WT mice, and the fate of these *Lbx1*+ cells may change.

Changes in interneuron identity have implications for reflex arcs, integrating complex afferent information, and signalling to higher brain centres (Bekkers et al., 2014). GABAergic neurons in the spinal cord act to ‘gate’ the strength of sensory input in primary sensory afferents and are critical in ‘normal’ function; preferences for excitatory networks have been implicated in chronic pain in humans (Takazawa and MacDermott, 2010; Zeilhofer et al., 2012), disrupted inhibitory networks have led to altered proprioceptive function in lamprey (Svensson et al., 2013).

### Impaired proprioception as a mechanism for AIS

The changes in vertebral stability, as demonstrated by increased rotation in the AIS-CRMΔ mice, with AIS-CRMΔ mice performing poorly on simple proprioceptive tasks, provide an initial possible mechanism for further investigation and further understanding of the role of proprioception in AIS. Although these findings are not as striking as those in the *Runx3* mutant described by Blecher et al. (2017), they also provide a plausible link between proprioceptive dysfunction and vertebral rotation.

Research has found that proprioception develops and functions from an early age, reaches maturity in adulthood, and subsequently declines with age (Martel et al., 2021; Viel et al., 2009). This study has shown that AIS-CRMΔ mice exhibit an early proprioceptive deficit, which precedes the onset of rotational deviation in the vertebral column. This deficit explains the vertebral deviation exhibited and follows previous studies showing the development of vertebral rotation and scoliosis upon loss of proprioceptive neurons (Blecher et al., 2017). Proprioceptive deficits have also been found between individuals with AIS and without AIS (Lau et al., 2022). While not evident in all populations, a similar mechanism involving other genes leading to proprioceptive deficits may contribute to AIS. It is notable that the current GWAS associating rs11190870 with AIS did not include proprioceptive testing in their studies because of the large cohort sizes (Assaiante et al., 2012; Lau et al., 2022).

The study did not show any notable differences between the sexes. The relationship between genetics and biological sex is intricate, with about 37% of genes showing some level of sex-specific expression in at least one tissue (Oliva et al., 2020). Additionally, environmental factors contribute to variability in the effects of sex hormones on individuals and populations. Although we investigated the influence of sex in our AIS-CRMΔ mice, we did not examine the individual effects of hormones in this context. To gain insight into sex-specific effects during puberty in the AIS-CRMΔ line, gonadectomy (orchiectomy or ovariectomy) could be performed on male and female mice before puberty.

### Future directions

Our study used whole tissue samples, which limits our ability to identify changes at the cellular level. Future investigations should use single-cell RNA-sequencing to examine changes in specific cell populations within different spinal cord regions during critical developmental periods to address this limitation. Analyzing these cell population differences may provide valuable insights into the cellular profiles and subsequent spinal cord functionality when AIS-CRM is deleted.

Our findings indicate that AIS-CRMΔ mice show increased vertebral rotation and poor performance on proprioceptive tasks, which suggests a possible mechanism that requires further investigation into the role of proprioception in AIS. Altered spinal circuitry, favoring increased GABAergic/inhibitory signaling, may have functional consequences such as chronic stimulation of associated paraspinal muscles and subsequent muscle atrophy, contributing to the phenotypic differences observed between WT and AIS-CRMΔ mice. Investigating the possible consequences of changes in *Lbx1* gene expression is also crucial as it regulates neuronal population numbers, migration, and cell fate, and its loss leads to changes in neurotransmitter identity, such as the switch from GABAergic to glutamatergic. Our study focused primarily on *Lbx1*, but other genes within the same TAD, such as *Fgf8, Poll*, *Btrc,* and *Fbxw4*, should be further investigated for changes in neuronal expression.

### Summary

Our findings suggest that AIS-CRM influences gene expression during a crucial embryonic period in the developing spinal cord (E10.5-E15.5). These effects were not limited to *Lbx1* but also extended to other genes such as *Poll, Fgf8*, *Fbxw4*, and *Btrc*. Together, these results suggest that the spinal cord contributes to AIS through changes in *Lbx1* and that the embryonic stages of neural tube development are the time point likely to influence spinal cord function postnatally. Further research into changes in neural tube patterning and specific neuron subtypes and how these persist postnatally remains to be investigated. Nevertheless, this is the first evidence to link the genetic variant of *Lbx1,* rs11190870, with the role of *Lbx1* in spinal cord development and subsequent functional deficits, resulting in a phenotype in mice that aligns with one aspect of the human phenotype of AIS.

## Acknowledgements

This research project was funded by a University of Otago Research Grant (UORG) to MJW. . JML was supported by the University of Otago’s postgraduate scholarship. We thank Andrew McNaughton from *Otago Micro and Nano Imaging Unit* who helped with establishing the microCT protocol.

## Methods

### Generation of the deletion line and Genotyping

To examine the effects of the human gene variant rs11190870 on spinal cord development and its possible contributions to AIS, a homologous, highly conserved region in the mouse was deleted using CRISPR-Cas9. A custom mouse line was obtained from Australian BioResources at the Garvan Institute (Australia). The deleted sequences are shown in Fig. S10. Two founder breeding pairs were sent, and the male and female founders were heterozygous. For genotyping, tissue samples were incubated in Cell/Tissue lysis buffer (10 mM NaCl, 10 mM Tris-HCl) with 20 mg/ml proteinase K at 55°C for a minimum of 2 h until the tissue was dissolved, and proteinase K was inactivated by incubating the samples at 85°C for 1 h. The samples were then spun at 14,000 g for 3 min to pellet any cellular/tissue debris, and the supernatant was transferred to a tube containing 500 μl of isopropanol and spun for 3 min to pellet the DNA. Finally, DNA pellets were resuspended in 300 μl of nuclease-free water. Polymerase chain reaction was performed using DreamTaq Hot Start Green DNA Polymerase (ThermoFisher), with an annealing temperature of 60°C. Oligonucleotide primer sequences can be found in Table S1. PCR products were analysed on a 3% agarose gel; the WT amplicon was 639 bp, the AIS-CRMΔ amplicon was 450 bp, and HET bands were present.

### In silico analysis of rs11190870

HiC and 4C data around SNP rs11190870 was accessed from http://3dgenome.fsm.northwestern.edu/. ENCODE data from www.encodeproject.org/. Data tracks were added to the UCSC genome browser for the mouse (mm10) and Human (hg19) for viewing concerning the target genome region. RegulomeDB https://regulomedb.org/ was additionally used for chromatin state analysis.

ReMap (2022) was loaded into the UCSC browser, this is a database of all public (Hammal et al., 2022). For biotypes, ChIP-seq data from mouse and human tissues included neuronal tissues, cell lines (NPC, NSC, ESC), and fetal tissues, were chosen. The following transcriptional regulators were selected: CTCF, EZH2, RAD21, SMC1, SCM3, SMC4, STAG1, and STAG2 (Cohesin proteins). The selection is limited to available data, and not all TFs selected have been analyzed for each tissue type.

### RNA collection from embryos

Following the euthanization of the dams, the uterus was placed in ice-cold 1x PBS. The neural tube was carefully dissected from the base of the developing brain vesicle to the caudal-most region, and alternative tissues were collected for genotyping. The samples were homogenized in 1 mL of TRIzol (Thermo Fisher, Cat. No. 15596026). Chloroform (200 μl) was added to the sample and incubated at R.T. 2 - 3 min before centrifugation. The aqueous layer was then transferred to a new RNase-free tube and combined with 500 μl of isopropanol to precipitate the RNA before spinning for 15 min at 4°C to pellet the RNA. The pellet was washed in ice-cold 70% ethanol twice, and the air-dried RNA pellet was resuspended in 50 μL of nuclease-free water. RNA samples were stored at −20°C for short-term storage or at −80°C for long-term storage. RNA concentration and purity were measured using Nanodrop2000 (Thermo Fisher Scientific). Samples with a 260:280 ratio between 1.90-2.10 were deemed acceptable for downstream applications.

### RT-qPCR

Following RNA extraction, complementary DNA (cDNA) was synthesized from a total of 1 μg RNA in a 20 μl reaction mix using the QuantaBio qScript XLT supermix (Cat. no. 95161-100) as per the manufacturer’s instructions. RT-qPCR was performed using a QuantStudio3 Real-Time PCR system (ThermoFisher). Reactions consisted of 5 μl PowerUp^TM^ SYBR^TM^ Green Master Mix (Applied Biosystems), 1 μl primer working stock at 20 pmol/μl (10 pmol forward, 10 pmol reverse) and 3 μl nuclease-free H_2_O, and 1 μl of cDNA. All reactions were run in triplicate per biological replicate. Relative expression was calculated using two reference genes, *Pgk1* and *Sdha*. Fold change in expression was calculated using the Livak method (2^−ΔΔCt^), relative to WT.

### MicroCT Imaging of the vertebral column

We used a post-mortem micro-CT scan to provide a detailed 3D view of the vertebral column. Previous studies have used plastic straws and similar objects to secure their spines in place; however, there was concern that a plastic straw or similar object would interfere with any observable phenotype. Therefore, a ‘stand’ was developed (Fig. S11). This ensured there was little movement to the spine during the setup and imaging; it provided a humid environment that stopped desiccation.

The SkyScan 1172 source voltage was set to 50 kV and the source current to 200uA with a 0.5 mm aluminium filter. Image pixel size (resolution) was set to 34.78μm with a 0.5-degree rotation step. Following imaging, 3D scans were reconstructed using NRecon (1.7.4.6), allowing for FIJI (ImageJ) analysis. The stand and other materials were filtered and excluded from the 3-dimensional reconstructions during reconstruction. Vertebral bodies were individually reconstructed from the source image, and the rotation angle was measured relative to the L5 vertebrae. Angular measurements were compared using the GraphPad Prism software.

### Behavioural Testing

Upon entry into the study, the animals were housed in the behavioural phenotyping unit at The University of Otago (BPU) for 2-3 days before their first testing day for acclimation. The animals were weighed and moved from their housing space at the BPU (University of Otago, N.Z.) to a testing room before testing. Baseline recordings of the behavioural tests were conducted to obtain the initial behavioural readings for each litter. The baseline grid walks, open field, and SNAP tests were carried out on postnatal day 28 (P28), while the baseline cylinder and grip strength tests were carried out two days later at P30. After obtaining baseline readings, behavioural testing was repeated every two weeks for two testing days for each litter. Mice were weighed before every testing session to monitor their health and for later use in behavioural analysis. Observers performing the behavioural analysis were blinded to the genotype of the mice.

### Adapting Simple Neuroassessment for Asymmetric Impairment (SNAP) testing to assess proprioceptive dysfunction

The Simple Neuroassement for Asymmetric Impairment (SNAP) is an observational assessment used to quantify neurological deficits in a mouse model of traumatic brain injury (TBI) mouse model (Shelton et al., 2008). It consists of seven tests to evaluate emotional and physical behaviours in mice. Shelton et al. (2008) observed proprioceptive deficits in their model of TBI; as such, this was adapted to score the observed proprioceptive defects in our cohort. A detailed description of the scoring metrics can be found in Table S3. Briefly, we assessed six behavioural tasks: general activity, interactions (with handlers), cage grasp, visual placing, gait/posture, and baton/grip test. Animals were scored from to 0-5, with 0 being ‘typical/no defect’ and 5 being severe impairment/deficit. The scores were summed, averaged across the cohorts, and plotted to provide an overall SNAP score for the mice comparing WT and AIS-CRMΔ littermates. All tests were performed blinded to the genotype. In addition, the data were analysed using two-way ANOVA to account for age and genotype.

### Grid Walk Test

The grid walk test assesses sensorimotor function by testing motor coordination and foot-placing deficits during locomotion (Russel et al., 2011). It is also a simple and effective way to assess proprioceptive dysfunction by comparing the number of foot faults to the total number of steps taken. Mice were placed in a plexiglass box on a gridded area (32 × 20 × 50 cm) with 11 × 11 mm diameter openings. Behaviour was recorded using a camera placed on a tripod facing down at a mirror beneath the grid to assess the animal’s stepping errors (i.e., foot faults). The animals were given 5 min to walk around the gridded area. The foot faults for each limb were counted and compared with the number of steps made by that limb. A step was considered a foot fault if it did not provide support and the foot slipped through the grid hole. The percentage of foot faults out of the total number of steps was calculated for each limb and used to compare sensorimotor function between the limbs and animals.

### Grip Strength Test

The grip strength test is often used to evaluate motor function and deficits in mice with CNS disorders such as proprioceptive dysfunction (Maurissen et al., 2003). This test was used to quantify the skeletal and muscular strength of the forelimb and hindlimbs in mice. The grip strength apparatus (Model BIO-GS3, Bioseb) comprised a bar connected to an isometric force transducer (dynamometer) for forelimb testing and a wire grid (8 × 8 cm). The force transducer meters were set to zero and reset before each measurement to allow for the detection of proper values. The force was measured in grams (g). After visually checking that the grip was symmetric and tight, the mice were gently pulled horizontally by the tail until the grasp of the bar was broken (Takeshita et al., 2017). To measure the combined limb grip strength, the mice were allowed to rest on the angled wire grid and were pulled by the tail until their grip was broken. A dynamometer reduces the value at the maximal force/grip (Takeshita et al., 2017). Each test was repeated three times to obtain the maximal forelimb and combined limb grip strength for each mouse. Hind limb grip strength was calculated by subtracting the maximal forelimb grip strength from the maximal combined grip strength. Body weight was measured before each testing session and used to normalise grip strength results (Takeshita et al., 2017).

### Cylinder Test

The cylinder test assessed CNS and proprioceptive function by measuring rearing frequency and paw preference. The animals were placed in an upright clear plexiglass cylinder (100 mm diameter × 150 mm height) with a mirror positioned behind the cylinder, allowing all movements to be captured. Videos (5 min) of animal activity were recorded and assessed for forelimb placement, slips, and other rearing behaviours. Mice often rear up to a standing position to explore and press on the cylinder wall with their left, right, or both forelimbs for support. The ratio of the time spent on the left forelimb relative to the right forelimb was also calculated to measure forelimb asymmetry.

### Immunofluorescence

Transverse cryosections of the neural tubes and adult spinal cords were collected and mounted directly onto slides. Slides were defrosted and allowed to come to room temperature (R.T.) to dry. Immunofluorescence for LBX1 was carried out as described in (Decourtye et al., 2022)

### ChIP-qPCR

Tissue for chromatin extraction was prepared from the embryos (E12.5). Brain and neural tubes were dissected from the embryos and cross-linked for 10 min with methanol-free formaldehyde (ThermoFisher #28906). The cross-linking reactions were stopped with 0.125 M glycine before being spun down, and the tissue pellet was frozen at −80°C for storage.

To prepare the beads, 50 <u>μ</u>l of protein G Dynabeads (ThermoFisher) was placed into a microfuge tube with 1 ml of block solution (0.5% bovine serum albumin in PBS). Beads were washed with block solution three times, re-suspended in 1.5 ml of block solution, and split between two microfuge tubes. The EZH2 antibody (10 mg) was added to one tube, and the other tube received IgG (pre-immune serum) as a mock control. Tubes were topped up to 1.5 ml with block solution, then rocked overnight at 4°C.

Following overnight incubation, 1ml of membrane extraction buffer with 1% proteinase inhibitor (ThermoFisher) was added, and tissue was broken up with an 18g syringe. The tissue was spun at 9000 x g for 3 min. The supernatant was removed, and tissue was resuspended in 500 μl of digestion buffer with 1 μl of 500 nM Dithiothreitol (DTT). The tissue was digested with 1ul MNase (ThermoFisher; 100 units) in 500 μl volume at 37°C for 15 min. 50 ml of stop solution was added, and the sample was left on ice for 5 min. This was centrifuged at 9000 × g for 3 min, and the supernatant was removed. Nuclear extraction buffer (250 mL) was added to 2.5 ml of proteinase inhibitor, and the sample was left on ice for 15 min and vortexed for 15 s every 5 min. The sample was spun at 9000 x g for 5 min, and the supernatant was collected. Fifty μl of supernatant was reserved for input.

Unbound antibody was removed by washing the beads with a block solution. The beads were then re-suspended in 1.25 ml of block solution. The chromatin solution was split equally between the beads, and the chromatin-antibody samples were rocked at 4°C overnight. Beads were washed four times with wash buffer 1 (50 mM HEPES-KOH pH 7.5, 500 mM LiCl, 1 mM EDTA, 1% Nonident P-40, 0.7% Sodium deoxycholate) for 15 min each, rotated between washes, and then rinsed in wash buffer 2 (TE pH 8.0, 50 mM NaCl) for 5 min. Elution buffer (100 µL) was added, and the samples were incubated at 65°C for 10 min. Chromatin (50 µL) was reserved for a input control during chromatin preparation. Samples were incubated with 5 M NaCl and Proteinase K at 65°C for 2 h and followed by phenol: chloroform precipitation and ethanol precipitation. The resulting pellets were dissolved in 50 µL of H_2_O.

ChIP-qPCR reactions were repeated in triplicate, and amplification of the region of interest was deemed successful if the melt curve produced a single peak and Ct values in triplicate were within < 0.5. The resulting Ct values were then used to calculate the percentage pulldown compared to the no antibody/mock control using the following formula: % pulldown = 100 × _2(Ct adjusted input - Ct antibody) /2(Ct adjusted input - Ct mock)._

### Statistical analysis

All statistical tests were performed using Prism 9. The effect of three variables (AIS-CRM genotype, age, and sex) was determined using three-way analysis of variance (ANOVA) with multiple comparisons (Tukey’s multiple comparisons test). To determine the effect of two variables, AIS-CRM genotype and age, a two-way ANOVA with multiple comparison testing was conducted. Pairwise comparisons were performed using unpaired t tests.

