## Supplementary Information for "Deletion of a conserved genomic region associated with adolescent idiopathic scoliosis leads to vertebral rotation in mice"

**Supplementary Files:**

File S1 RegulomeDB analyses output.

File S2: 3-way ANOVA testing results.

**Table S1. Oligonucleotide sequences (5’-3’).**

| | Forward | Reverse |
|---|---|---|
| Genotyping primers | | |
| *AIS-CRM* | GCTGTGGCTTGAGTCGTTATC | TGTTTTCCCTTTAAATCGAGGAC |
| *Sry* | TCATGAGACTGCCAACCACAG | CATGACCACCACCACCACCAA |
| qPCR primers | | |
| *Pgk1* | CACAGAAGGCTGGTGGATTT | CTTTAGCGCCTCCCAAGATAG |
| *Sdha* | GCTGGAGAAGAATCGGTTATGA | GCATCGACTTCTGCATGTTTAG |
| *Btrc* | TGTGGCCAAAACAAAACTTGCC | ATCTGACTCTGACCACTGCT |
| *Dpcd* | CAAAGAAAGTAATGCCAATCCCA | TTCGAATCCGCCACTGGAAA |
| *Lbx1* | GCGATGGGATGACCATCTTT | GCGTTTCTCCAACTCGTAGAT |
| *Npm3* | AGGCAGGGTACAGCAAGTTC | CCCTCTCCACTCCCGACTTA |
| *Fbxw4* | GACAGCCGGCTTTAGGTACA | GTTCTGCCCAGTGTATGCCT |
| Poll | AGAGCCCGCTCATAGTCCA | CCGGGCTGAACTCTTTGAGAA |

#### Table S2: Genes located within the LBX1 TAD.

| Gene | Product | Function | Expression | Associated Diseases |
|---|---|---|---|---|
| *Poll* | DNA Polymerase Lambda[37] | - DNA repair, base excision repair[38], NHEJ[37]<br>- Roles in meiosis[39] | - Ubiquitous expression[39]<br>- Testes[39] | - Deletion of *Poll* results in disruption to *Dpcd*<br>- Loss of ciliated cells resulting in pulmonary dysfunction<br>- *Situs inversus*[40]<br>- Split-hand/foot |
| *Btrc* | F-box protein subunit[42] | - Mediates cell cycle progression[43]<br>- F-box protein mediates ubiquitination/degradation of β-Catenin[44] | - Ubiquitous | - Split-hand/foot malformation 3[41] |
| *Dpcd* | Protein involved in the generation of ciliated cells | - Expression increases during ciliated cell differentiation[45]<br>- Disruption results in primary ciliary diskenesia[45] | - Airway epithelial cells[45,46]<br>- Ciliated cells[45,46]<br>- Fagella-containing cells[45,46] | - Loss of ciliated cells resulting in pulmonary dysfunction<br>- *Situs inversus*[40] |
| *Fgf8* | Protein belonging to Fgf family | - Important regulator of embryonic development, cell proliferation, differentiation and migration[47–50].<br>- Left/Right axis determination[47]<br>- Normal development of GnRH neuronal system[50] | - Development: eye, brain, limb and ear[47–49] | - Hypogonadotrophic Hypogonadism 6<br>- Split-hand/foot malformation 3[41] |
| *Fbxw4* | Member of F-box/WD40 gene family | - Mediates degradation of target genes through ubiquitination[51]<br>- Potentially maintenance of AER[49] | - Apical Ectodermal Ridge/Limb[49] | - Split-hand/foot malformation 3[41] |
| *Npm3* | Nuclear chaperone protein | - Nuclear Chaperone[52]<br>- Chromatin remodelling[53] | - Ubiquitous | - Lung Papillary adenocarcinoma |

**Table S3. SNAP Scoring sheet.** Adapted from (Shelton et al., 2008)

| Behaviour | Score |
|---|---|
| Does the mouse look active? Is it placid/ does it move much when the bedding/<br>enrichment is removed? | 0. Mouse behaves normally and runs around cage.<br>2. Mouse movement is sporadic, often takes pauses.<br>5. Mouse doesn't appear active. |
| Is the mouse alert and avoiding the handler, are you able to easily touch the mouse? | 0. Mouse behaves normally and evades handler.<br>2. Mouse moves after handler gently nudges or touches mouse.<br>5. Mouse doesn't move even when touched by handler. |
| Taking the mouse by the base of the tail, lowering it to where it can grasp the metal bars.<br>on the food catcher, is the strength stronger/weaker than its WT littermate? | • Is there a particular paw that isn't grasping?<br>• Is it easy to pull the mouse away from the cage?<br>0. Mouse finds metal bar quickly and grabs onto it with both paws. Strong grip before being pulled away.<br>2. Mouse finds metal bar but let's go with one hand or is weaker than matched control.<br>5. Mouse struggles to grip on to the metal bar, very weak grip, let's go with both hands. frequently. |
| Taking the mouse by the base of the tail, when lifting the mouse towards a ledge, does it lift its head and extend its forelimbs to move | 0. Mouse finds ledge quickly and grabs onto it with both paws.<br>2. Mouse twists away from ledge before finally finding/gripping it.<br>5. Mouse doesn't find ledge or grab onto it after several failed attempts. |
| Where are its limbs positioned, under the body, out the side? | • Mice with proprioceptive defect will have limbs, particularly hindlimbs, splayed out<br>the side when walking on flat surface.<br>• How does its walk compare to littermates?<br>Score as below:<br>0. Mouse walks normally with all limbs positioned under its body.<br>2. Mouse appears to walk normally but limbs are slightly away from midline.<br>5. Mouse appears to have 'wobbly' gait, limbs are splayed and not kept close to midline. |
| Holding the mouse by its tail, place a long-handled applicator (cotton tip, or other similar<br>thickness stick) close to the mouse and observe its grip | • Does it grip with all four paws?<br>• How quickly does it realise it can be gripped with its hind limbs? If you move the baton<br>towards/away from the mouse does it then grip the baton?<br>• Does it try to climb the baton?<br>Score as below:<br>0. Mouse grabs baton immediately with all four paws, will try to move up/down the baton.<br>2. Mouse grabs baton with either front or hindlimbs but not both, required encouragement to<br>hold baton.<br>5. Mouse fails to hold baton, even after encouragement. |

### Supplementary Figures

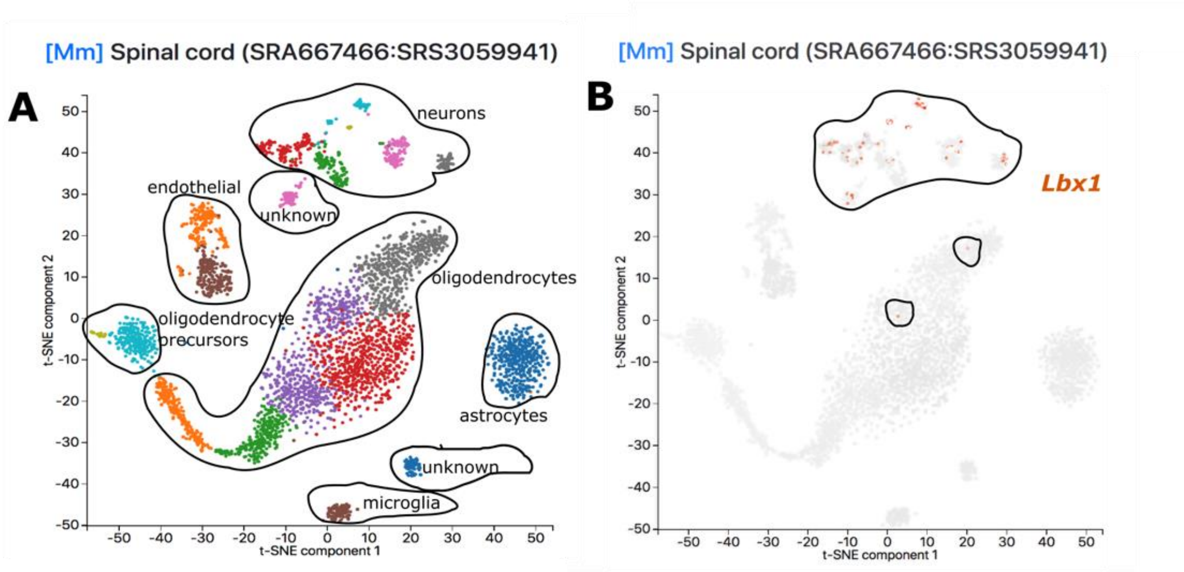


**Figure S1 Single-Cell Transcriptomic Analysis of the Adult Mouse Spinal Cord.** A) Clustering of cell types within the spinal cord of the adult mouse, adapted from PanglaoDB. B) *Lbx1* transcripts (orange cells) are restricted to the neuronal clusters with a few hits in the oligodendrocyte cluster. Data was sourced and adapted from PanglaoDB (https://panglaodb.se/) using the Spinal cord SRA667466:SRS3059941 dataset.

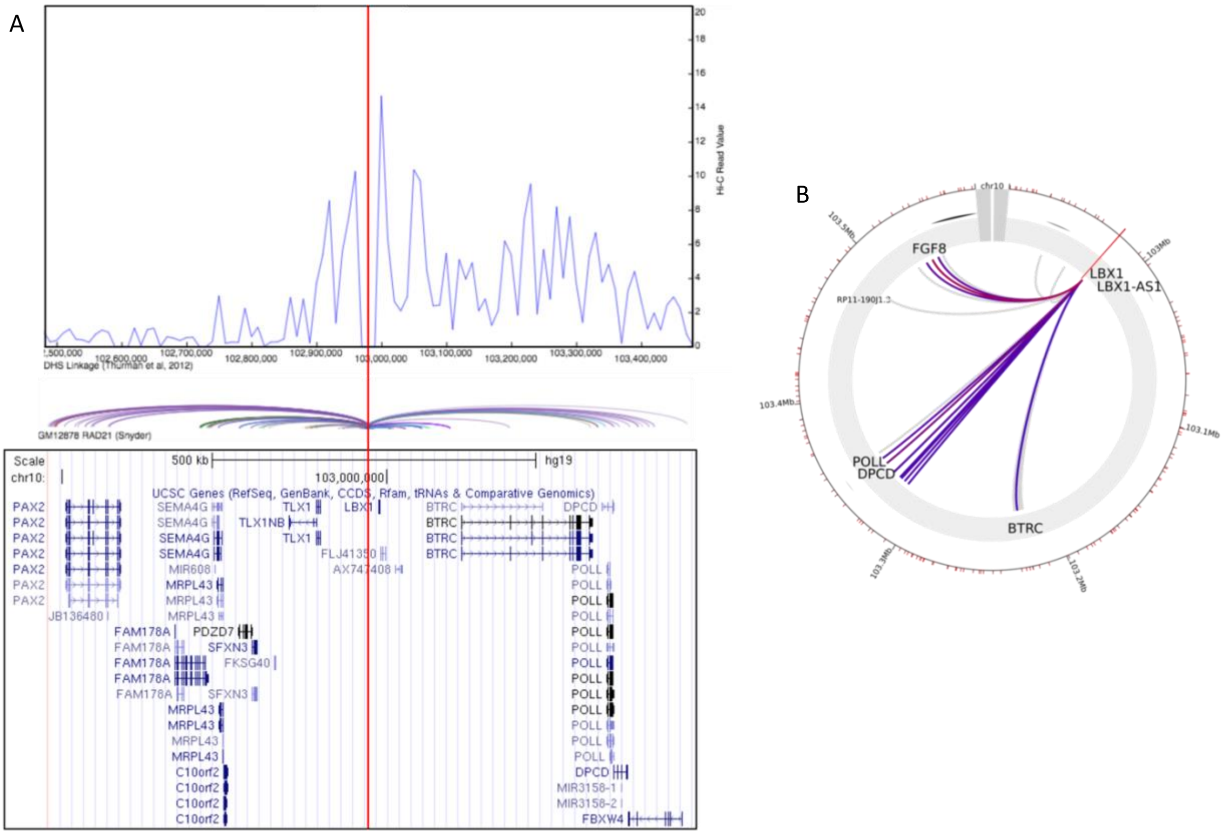


**Figure 1. Bioinformatic analysis of 4C/HiC data reveals possible contact of the putative enhancer with Lbx1 and genes surrounding Lbx1, including Poll, Btrc, Dpcd and Fgf8.** A) HiC contact map, red line indicates rs11190870 and is the anchoring point, while the blue lines represent the HiC read value. Generated using H1- Embryonic Stem Cells. B) Circos plot demonstrating rs11190870 contact with neighbouring genes and their respective distances generated using hESC.

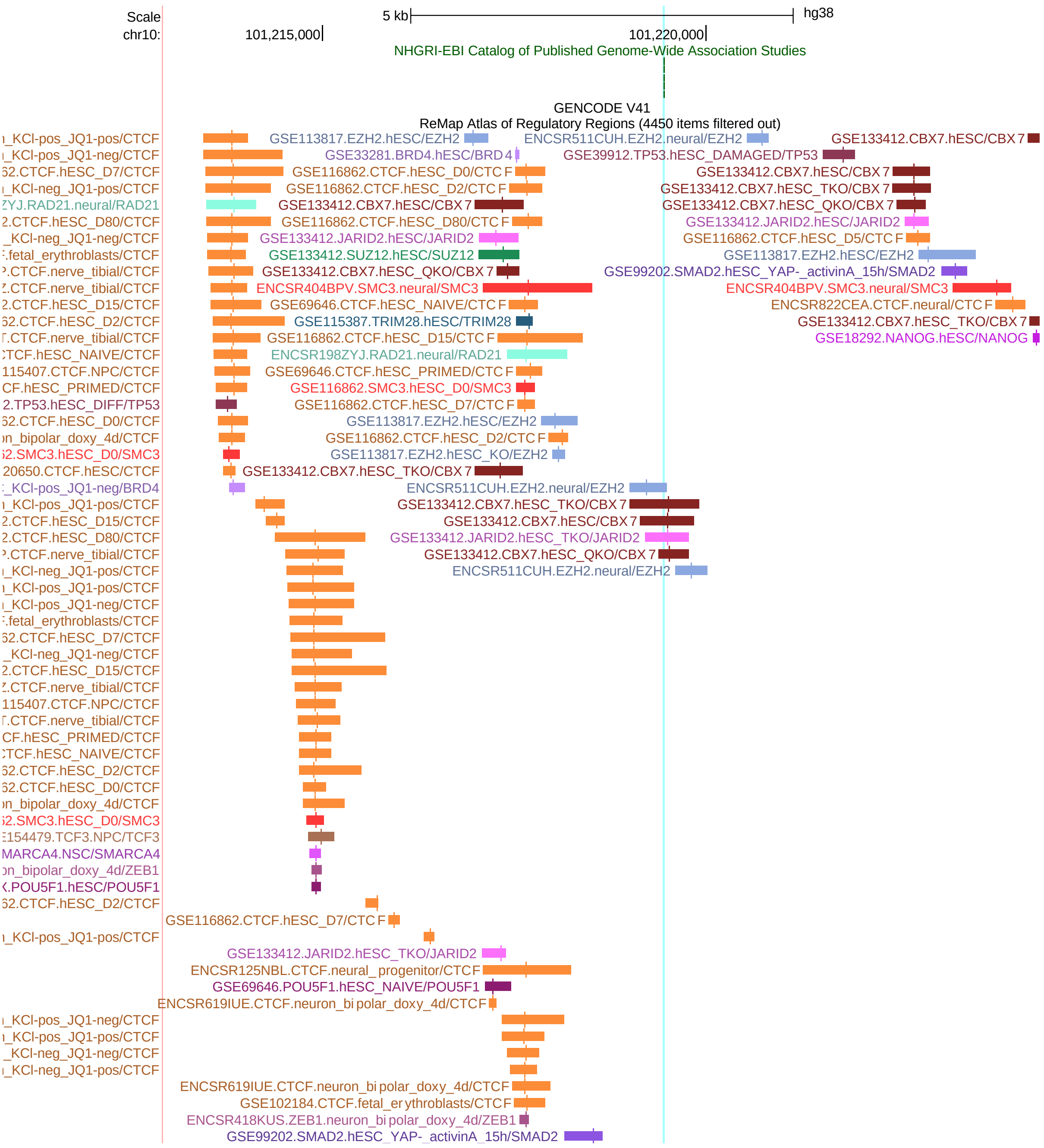


**Figure S3. Genome browser view (hg38)( RepMap2022) displaying ChIP-seq peaks for transcriptional regulators generated with neuronal samples**. The long vertical line (pale green) indicates the AIS-linked SNP. This area is consistent with a TAD border, marked by CTCF binding sites and Cohesin proteins. Each transcriptional regulator is colour coded, and the description includes the biological sample source for the ChIP experiment.

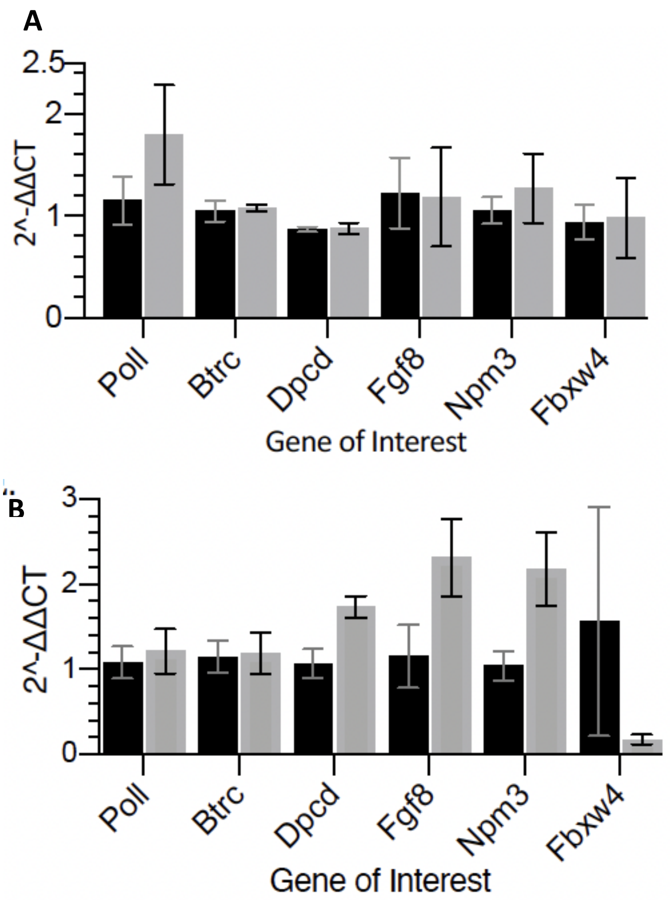


**Figure S4: Post-natal expression of genes associated with the TAD**. Transcript expression for *Btrc, Dpcd, Poll* and *Fgf8* in the KO relative to the wildtype cohort. Pre-Puberty (A) and Adult (B). Data is presented as mean SEM. Statistical significance was determined using 2way ANOVA, significance. Pre-puberty n=9 (WT), n=8 (KO); Adult n=7(WT), n=6 ( KO).

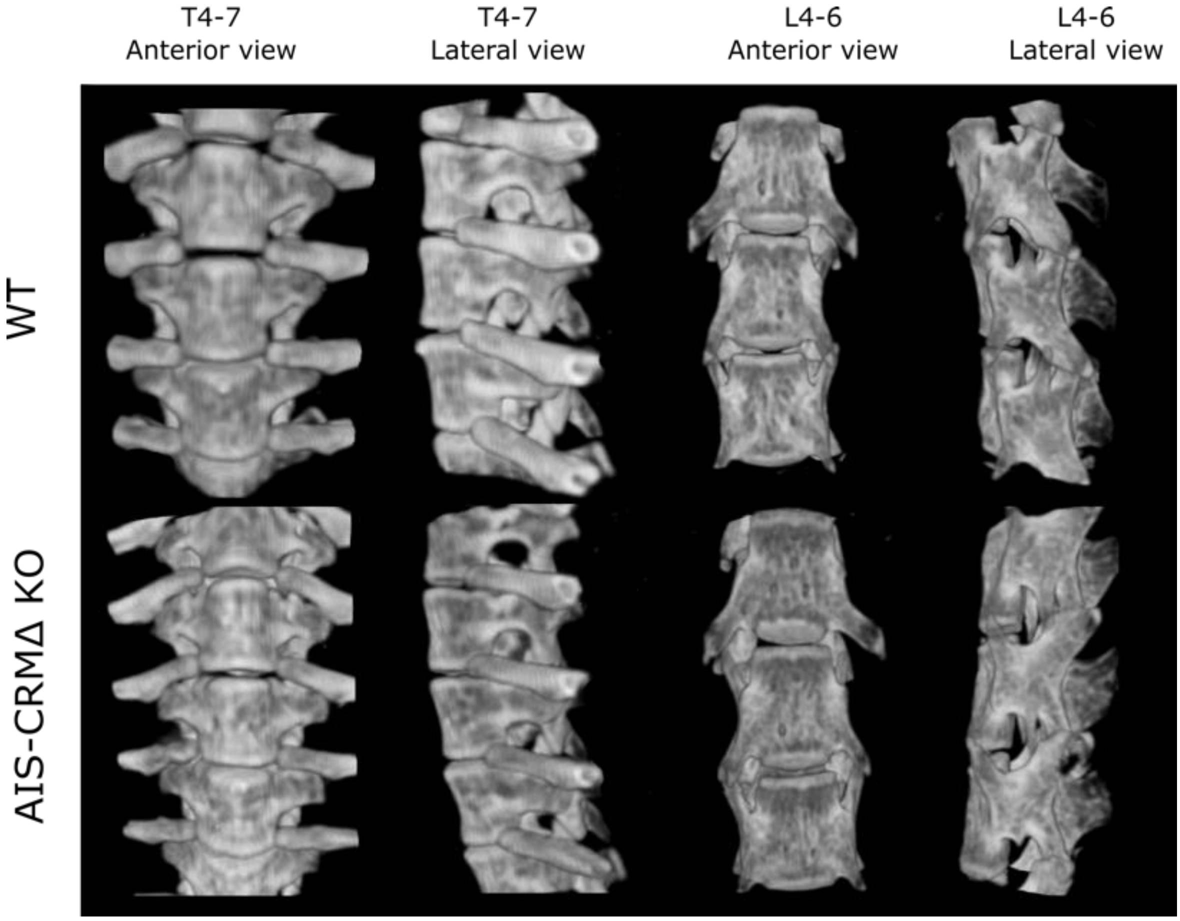


**Figure S5. Gross morphology and vertebral rotation in adult WT and AIS-CRMΔ mice using μCT.** Renderings of the vertebrae from both WT and AIS-CRMΔ mice revealed no gross structural difference between the vertebrae at all levels. Abbreviations: Thoracic (T), Lumbar (L).

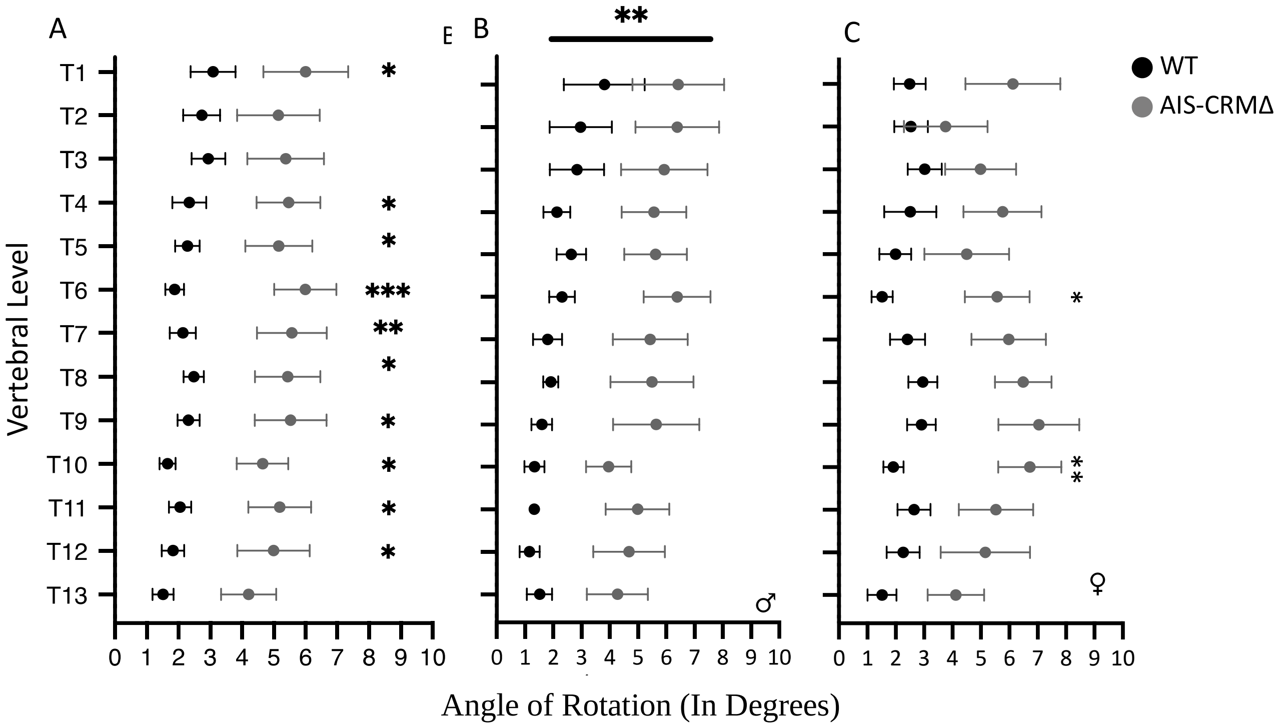


**Figure S6. The vertebral angle of rotation from WT and AIS-CRMΔ KO mice, the thoracic region only, segregated by sex.** (A) Degree of rotation relative to L5 in the thoracic region (T1-T13), combined sexes, (B) Male only and (C) Female only. Wildtype mice were represented in black (Combined n= 20, Male n= 9, Female n=11) and knockout in grey (Combined n=10, Male n=6, Female n=4). Measurements are presented as mean rotational deviation relative to L5. Data are presented as mean ±SEM. Statistical significance is represented as p<0.05=*, p<0.01=**, p<0.001=*** using a 2-Way ANOVA.

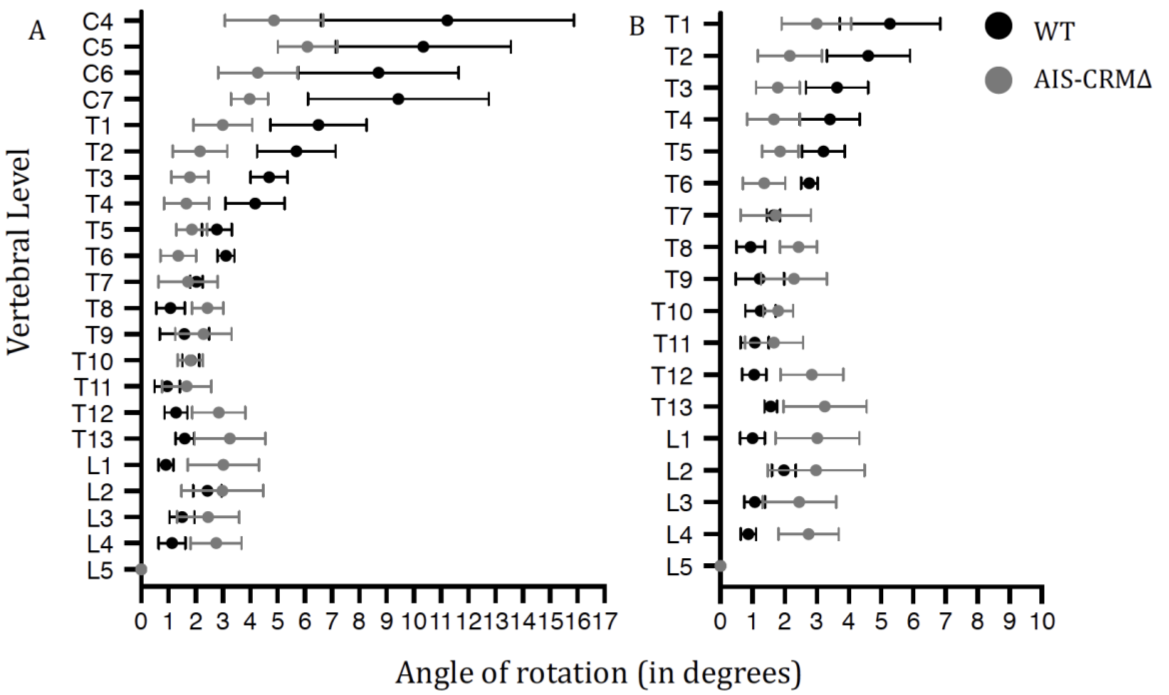


**Figure S7 Vertebral rotation angle in pre-puberty WT and AIS-CRMΔ mice using microCT.** Degree of rotation measured across C4 - L5 in the mouse samples (A), combining sexes. Wildtype mouse measurements are represented in black (n = 4) and knockout is represented in grey (n = 4). Measurements were then restricted to the thoracolumbar region (B), the region more commonly affected by AIS. Measurements are presented as a mean rotational deviation; error bars represent ±SEM. Statistical significance was calculated using 2-way ANOVA.

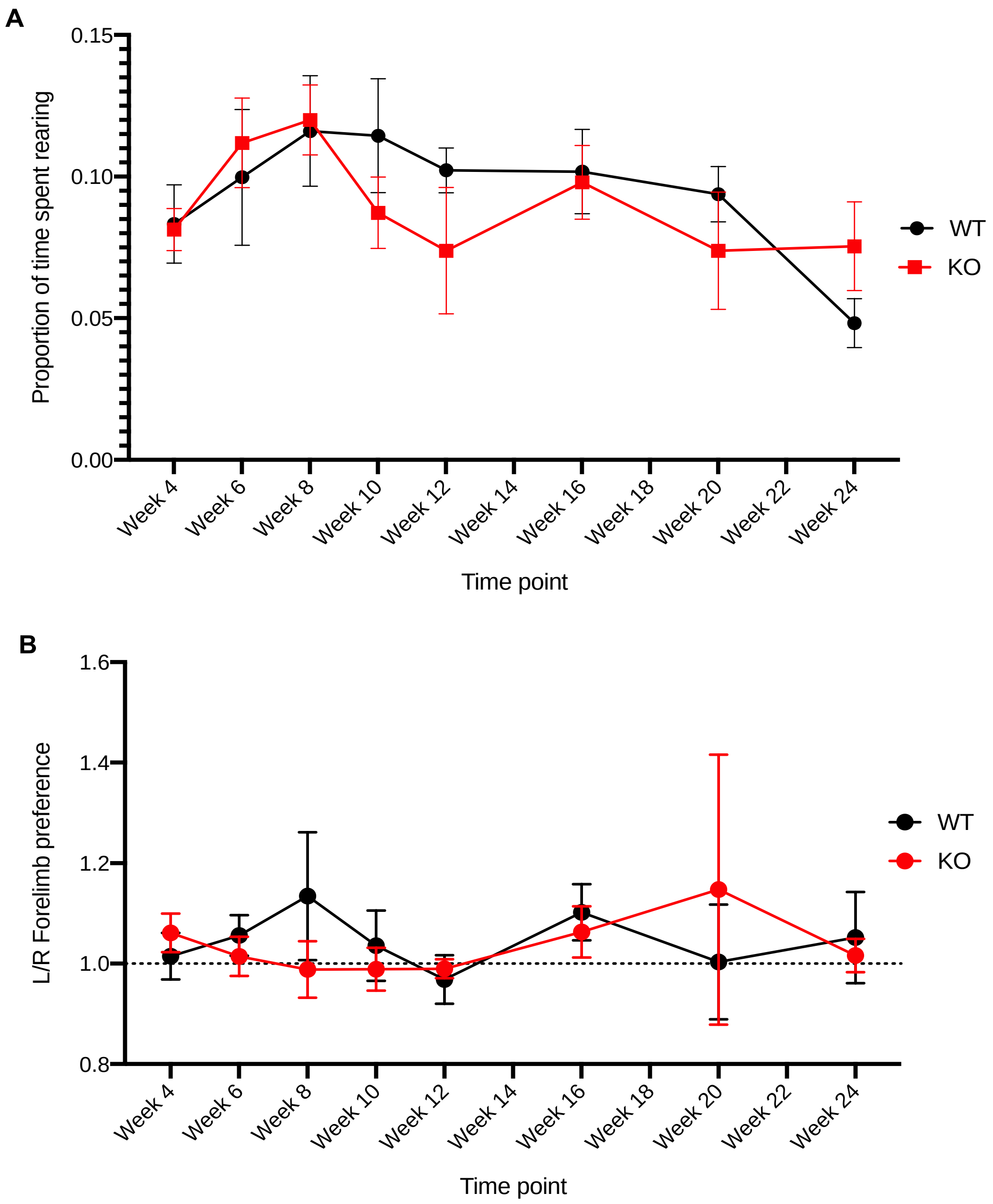


**Figure S8. The cylinder test measures the effect of the Lbx1EHΔ$^{-/-}$ deletion on motor function and motor asymmetry over 20 weeks.** Mice numbers assayed vary with each timepoint due to the timing of entry into the experiment; at least four WT mice (n = 4-8) and four KO (n = 4-13) were assayed. **A**. Difference in proportion of time subjects spent rearing and touching the cylinder wall between WT and Lbx1EHΔ$^{-/-}$ mice. **B**. Forelimb preference for animals when rearing and touching the cylinder wall over 20 weeks. Values over 1.0 represent left forelimb preference, while values below 1.0 represent right paw preference. All data were expressed as the mean ± SEM, where statistical analysis using a two-way ANOVA was defined as *$p<0.05$. Abbreviations: WT = Wild type, KO = Knockout.

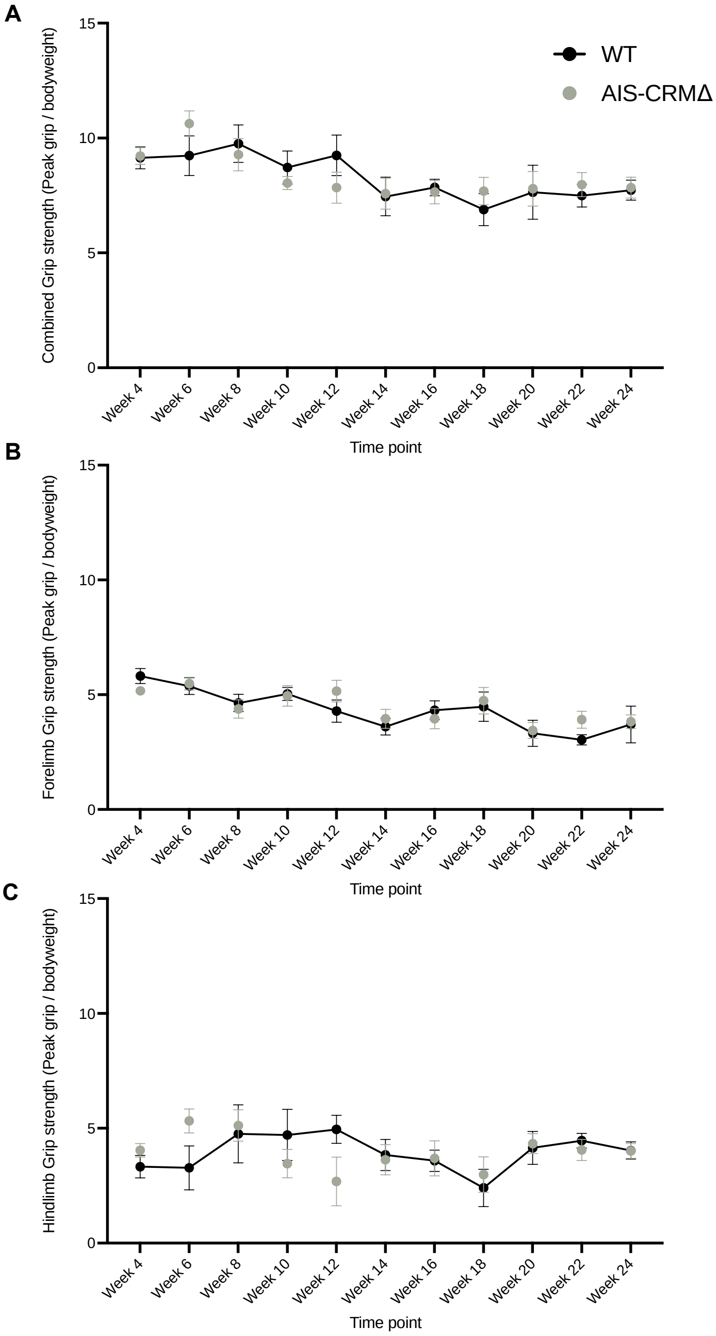


**Figure S9. Comparison of the effect of the AISCRMΔ on limb grip strength over 20 weeks.** Mice numbers assayed vary with each timepoint due to the timing of entry into the experiment; at least four WT mice (n=4-9) and four AIS-CRM (n=5-10) were assayed. **A)** Difference in forelimb grip strength between WT and AIS-CRM. A statistically significant decrease in forelimb grip strength from week 6 to week 8 and week 12 to week 24 was observed in the AIS-CRM A statistically significant decrease in forelimb grip strength was also observed in the WTs from week 4 to week 24. **B)** Difference in hindlimb grip strength between WT and AIS-CRMΔ. A statistically significant decrease in hindlimb grip strength was observed from week 6 to week 12 in the AIS-CRMΔs and from week 12 to week 18 in the WTs. All data are expressed as the mean ± SEM where statistical analysis using a two-way ANOVA was represented as **p<0.01. Abbreviations: WT = Wild type.

```
ttttagtcctggcgtttctgcagagcctcttaatacttcaaacttcaattttacgggcga
||||||||||||||||||||||||||||||||||||||||||||||||||||||||||||
ttttagtcctggcgtttctgcagagcctcttaatacttcaaacttcaattttacgggcga

ttgaactaagcatgtttctga---------------------------------------
|||||||||||||||||||||
ttgaactaagcatgtttctgacaaaattgatatatgtattaatttgctttaactggtgat

------------------------------------------------------------

taattactctccactgaaagaattatatggagctgtttgcctgcgatttgcatattaatc

------------------------------------------------------------

aaattctagttgataattctgccagtggggggggggtcagggctgggcacggggcacag

------------------------------tgtgtgccggtgtatgtgggggctccc
                              |||||||||||||||||||||||||||
ggtgtgaaggggcgcttgtgtgagtgccggtgtgtgccggtgtatgtgggggctccc

cacccccctcctttgctcgctccctgcttagctgccagtttagtgcattaaaaccaaagat
||||||||||||||||||||||||||||||||||||||||||||||||||||||||||||
cacccccctcctttgctcgctccctgcttagctgccagtttagtgcattaaaaccaaagat
```

**Figure S10. Genomic sequence deleted in the mouse line. Bottom strand is the WT sequence.** Two guide sequences (in green) to remove the targeted sequence (red).

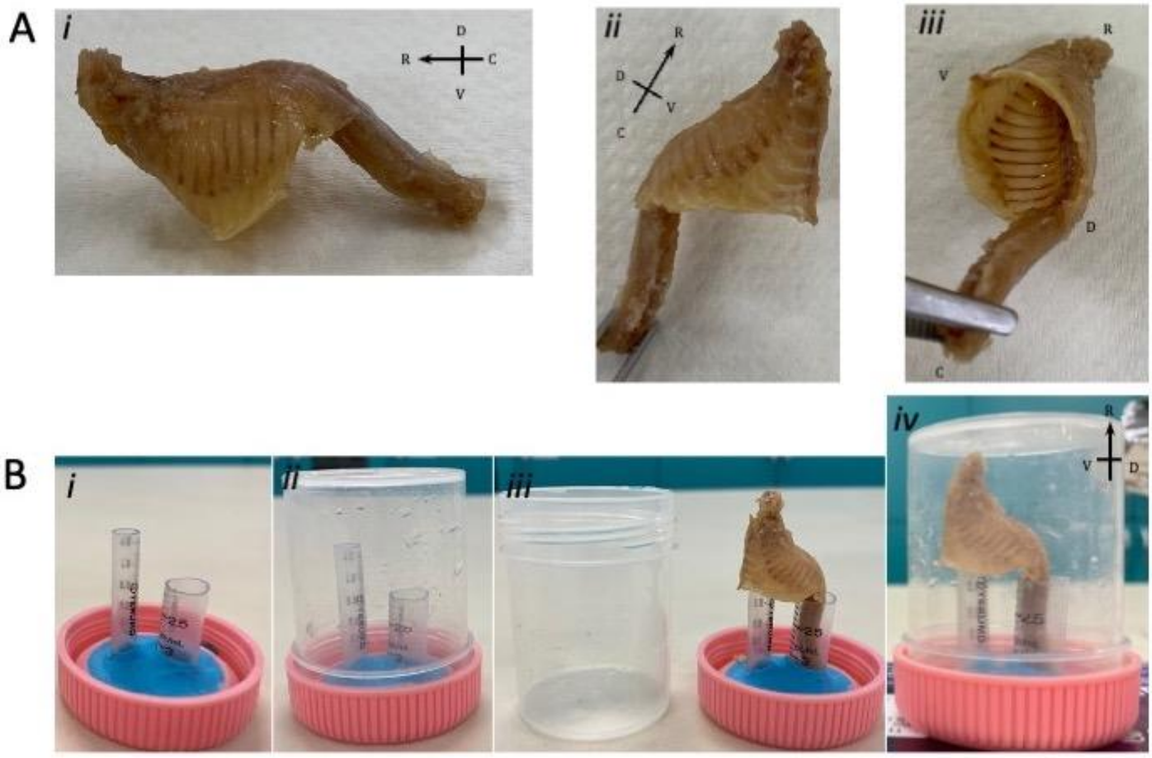


**Figure S11 Sample preparation for examining vertebral integrity and rotation in the AIS-CRMΔ mouse line.** Following euthanasia using by $CO_2$, mice are dissected, eviscerated, removing the head, limbs, and tail, and excess adipose/muscle is trimmed away. The samples should contain vertebrae C2-L7 and the thoracic cage. **Ai.** Lateral view of the sample following dissection. **Aii.** Lateral view of the sample following dissection. **Aiii.** Inferior view into thoracic cavity following evisceration. Spines are fixed overnight in 4% PFA (roughly 10x sample volume). Samples are trimmed of any remaining excess tissue. The sample is test fitted in the specimen pottle (**Bi-ii**) to ensure no contact between the pottle and specimen (**Biii-iv**). The specimen pottle was used to prevent desiccation and movement of the sample during the long scanning process. Samples undergo low-medium resolution CT scanning on the SkyScan 1172.
